# Mutual information estimation for transcriptional regulatory network inference

**DOI:** 10.1101/132647

**Authors:** Jonathan Ish-Horowicz, John Reid

## Abstract

Mutual information-based network inference algorithms are an important tool in the reverse-engineering of transcriptional regulatory networks, but all rely on estimates of the mutual information between the expression of pairs of genes. Various methods exist to compute estimates of the mutual information, but none have been firmly established as optimal for network inference. The performance of 9 mutual information estimation methods are compared using three popular network inference algorithms: CLR, MRNET and ARACNE. The performance of the estimators is compared on one synthetic and two real datasets. For estimators that discretise data, the effect of discretisation parameters are also studied in detail. Implementations of 5 estimators are provided in parallelised C++ with an R interface. These are faster than alternative implementations, with reductions in computation time up to a factor of 3,500.

**Results:** The B-spline estimator consistently performs well on real and synthetic datasets. CLR was found to be the best performing inference algorithm, corroborating previous results indicating that it is the state of the art mutual inference algorithm. It is also found to be robust to the mutual information estimation method and their parameters. Furthermore, when using an estimator that discretises expression data, using *N*^1*/*3^ bins for *N* samples gives the most accurate inferred network. This contradicts previous findings that suggested using *N* ^1*/*2^ bins.

## 1 Introduction

### 1.1 Background

Network inference is the reverse-engineering of transcriptional regulatory networks from high-throughput expression data. Transcriptional regulatory networks are the first level of a hierarchy of regulatory mechanisms that operate in cells and control gene expression. Proteins called transcription factors either activate or repress the transcription of a gene. Since these transcription factors are themselves the products of the expression of another gene, dependence between the expression of two genes suggests a regulatory link between them. This is the central assumption of network inference: that dependence between the expression of two genes is due to a functional relationship.

Network inference is a highly underdetermined problem. There are approximately 4,500 genes in the popular model organism *Escherichia coli* (*E. coli*). This corresponds to over 10 million pairwise interactions which must be estimated from samples that typically number less than 1000. Several unsupervised approaches have been developed to tackle this problem, with different classes of methods proving effective at predicting certain network motifs. It has been shown that combining predictions from different classes of network inference algorithm significantly improves the accuracy of inferred networks [1]. Examples of the techniques utilised by network inference algorithms include feature selection [2] and Bayesian networks [3]. Another class of methods utilise mutual information, an information-theoretic quantity that captures dependence between two random variables. As well as network inference, mutual information is used in biology as a distance measure for gene expression clustering [4]. It is often preferred to linear correlation measures due to its ability to capture nonlinear dependencies.

Mutual information-based network inference algorithms have several advantages. Specifically, they are well-suited to the direct, model-free inference of large-scale regulatory networks. Since they only consider pairwise interactions they are computationally affordable even for large numbers of genes [5]. Furthermore, considering pairwise interactions involves estimating bivariate probability distributions, meaning that they perform relatively well with low numbers of samples [6, 7]. However, mutual information-based methods are unable to predict the type or direction of a regulatory relationship, unlike feature selection-based methods.

### 1.2 Motivation and Aims

Estimating mutual information is known to be difficult [8] and various methods exist for computing an estimate. This study examines the effect of mutual information estimators on the accuracy of the inferred network. Three mutual information-based network inference algorithms, CLR [9], MRNET [10] and ARACNE [6] are used to infer the network from a matrix of gene expression values. These algorithms produce a score for each gene pair, the ranking of which indicates the confidence of an regulatory link between the two genes. A single network is then produced by thresholding these scores. This score is derived from the mutual information between the expression profiles of every gene pair, and so the method used to estimate mutual information can have a significant effect on the accuracy of the inferred network. Further details on the network inference algorithms can be found in Section S2.

CLR was the best-performing mutual information-based inference algorithm in the DREAM5 Network Inference Challenge [1] and has been used in the development of antibiotics [11] and microbial fuel cells [12]. ARACNE is another popular method that has been used in the study of brain tumours [13] and the sequencing of the wheat genome [14]. MRNET has been shown to be competitive with CLR and ARACNE [15].

This study compares the performance of CLR, MRNET and ARACNE when used with nine of the most popular mutual information estimators. These are the Maximum Likelihood, Miller-Madow, Chao-Shen, Shrinkage, B-spline, Kernel Density, *k*-nearest-neighbour, Pearson correlation and Spearman correlation estimators. Of these estimators, the first four are histogram-based and hence require data to discretised/binned. This study also investigates the use of the Bayesian Blocks algorithm in network inference. Bayesian Blocks was developed by Scargle et al. to model astronomical time series data using piecewise constant functions [16] and can be used to discretise continuous data into bins whose number and width are chosen to optimally fit data. The increased sophistication of Bayesian Blocks over simpler alternatives such as equal width bins should lead to more accurate estimates of probability distributions. It has previously been applied to network inference and was found to improve the accuracy of inferred networks [17].

**Table 1:** The properties of the datasets used in study [1]. For each dataset a subset of the genes are designated as potential regulators and the accuracy of inferred networks are only evaluated using the interactions of those genes. *Proportion of links* refers to the fraction of this subset of interactions that are regulatory links in the gold standard network. For the real datasets this is likely to be an underestimate. These values demonstrate the large class imbalance that is typical for network inference. For more detailed information on the data used in this work see Section S3.

| *Type* | *Organism* | *Genes* | *Regulators* | *Samples* | *Proportion of links* |
| --- | --- | --- | --- | --- | --- |
| Synthetic | n/a | 1,643 | 193 | 805 | 0.014 |
| Real | *E. coli* | 4,297 | 296 | 805 | 0.002 |
| Real | *S. cerevisiae* | 5,950 | 183 | 563 | 0.004 |

The estimators will be compared using one synthetic and two real datasets from the DREAM5 Network Inference Challenge [1]. The synthetic dataset was generated with GeneNetWeaver [18] and is labelled as *in silico*, while the real datasets are from *E. coli* and *Saccharomyces cerevisiae* (*S. cerevisiae*). Further details on the datasets are given in Table 1 and Section S3.

### 1.3 Structure and contributions

The following subsection reviews existing studies on this topic. Section 2 outlines the process of network inference, the mathematical definition of mutual information and discusses methods of mutual information estimation. Section 3 compares the performance of these mutual information estimators for network inference using datasets from the DREAM5 Network Inference Challenge [1]. Detailed descriptions of the mutual information estimators, the network inference algorithms and the performance metrics used in this and other studies on this topic can be found in the Supplementary Material.

The contributions of this work are as follows:

- This is the first evaluation of mutual information estimation using the three most popular mutual information-based network inference algorithms.
- This is the first comprehensive evaluation of the parameters of the Maximum Likelihood, Miller-Madow, Chao-Shen and Shrinkage estimators to include Bayesian Blocks.
- This is the first evaluation of mutual information estimation for network inference to use the area under precision-recall curve (AUPRC) as the evaluation metric, as is recommended for the evaluation of classification problems with a large class imbalance [19–21].
- This is the first systematic evaluation of the effect of Bayesian Blocks in network inference.
- Implementations of 5 estimators are provided in parallelised C++ with an R interface. These are faster than alternative implementations, with reductions in computation time up to a factor of 3,500.

### 1.4 Review of existing studies

Several existing studies examine mutual information estimation for network inference. However, the inference algorithms, mutual information estimators and performance metrics vary between them. The most recent study by Kurt et al. examined the performance of 11 mutual information estimators, including all 9 estimators used in this study [22]. This study included raw Pearson and Spearman correlation values as the additional estimators. For the histogram-based estimators, equal width and equal frequency binning was used. This study used the network inference algorithms RELNET, ARACNE and C3NET. The approaches were evaluated using two synthetic datasets generated by SynTRen [23], an alternative tool to GeneNetWeaver, and two real datasets. The real datasets were from *E. coli* and *S. cerevisiae*. The authors found that the B-spline estimator and those based on the linear Pearson and correlation led to the most accurate inferred network on all the datasets.

The C3NET algorithm [24] does not produce a ranking of scores for each gene pair. Instead, it uses a heuristic approach to produce a single network. It is therefore not possible to evaluate its predictions using the metric used in this study, AUPRC. Instead the authors use the F-score. The evaluation metrics used in this study and others are discussed in Section S4. The RELNET (Relevance Network) algorithm ranks gene pairs by the mutual information between their expressions, with a higher rank corresponding to a more confident prediction of an edge [25]. It had the lowest prediction accuracy for all mutual information estimators and so is not included in this study. For reasons why this simplistic approach is not effective see Section S2.

A study by Olsen et al. [7] compared five mutual information estimators (Maximum Likelihood, Miller-Madow, Shrinkage, Pearson correlation and Spearman correlation) with CLR, MRNET and ARACNE. The evaluation metrics were F-scores and areas under receiver operating characteristic (ROC) curve. The authors report that CLR and MRNET outperform ARACNE, and that the Pearson and Spearman correlation estimators perform best on real data. In addition, the Miller-Madow estimator performs well on synthetic data. Olsen et al. also find that equal frequency binning is preferable to equal width binning.

A comprehensive review of the existing literature on mutual information estimation is was completed by Walters-Williams and Li [26]. While this is not directly applied to network inference it is still a valuable survey and reports that the *k*-NN and Kernel Density estimators outperform the Maximum Likelihood estimator. They also report that the B-spline estimator provides equivalent performance to the Kernel Density and *k*-nearest-neighbour estimators in a much smaller computation time.

## 2 Methods

### 2.1 Problem formulation and workflow

Starting with a matrix of gene expression values *X ∈ ℛ^N×p^*, the aim of network inference is to infer a graph *𝒢* = (*V, E*), where each node *g ∈ V* represents a single gene and each edge *e ∈ E*, represents a regulatory link between two genes.

From *X*, a mutual information estimator computes a symmetric matrix of mutual information values of all gene pairs, whose *ij*-th element is the mutual information between columns *i* and *j* of *X*. The network inference algorithm then computes a score for each gene pair from these mutual information values. These scores reflect the confidence that there is an edge between two genes. To produce a single inferred network from these edge scores we choose a threshold and consider all scores above that threshold as edges, with all scores below designated as non-edges. This means that the scores themselves are not analysed, only their ranks. The inferred network is then evaluated against the true network and the precision and recall are calculated (see Section S4 for more details). This procedure is illustrated in Figure 1. This is typically repeated for all possible thresholds, resulting in a list of precision-recall values. These values are plotted and the area underneath the resulting curve is used to quantify the accuracy of the predictions.

**Figure 1:**
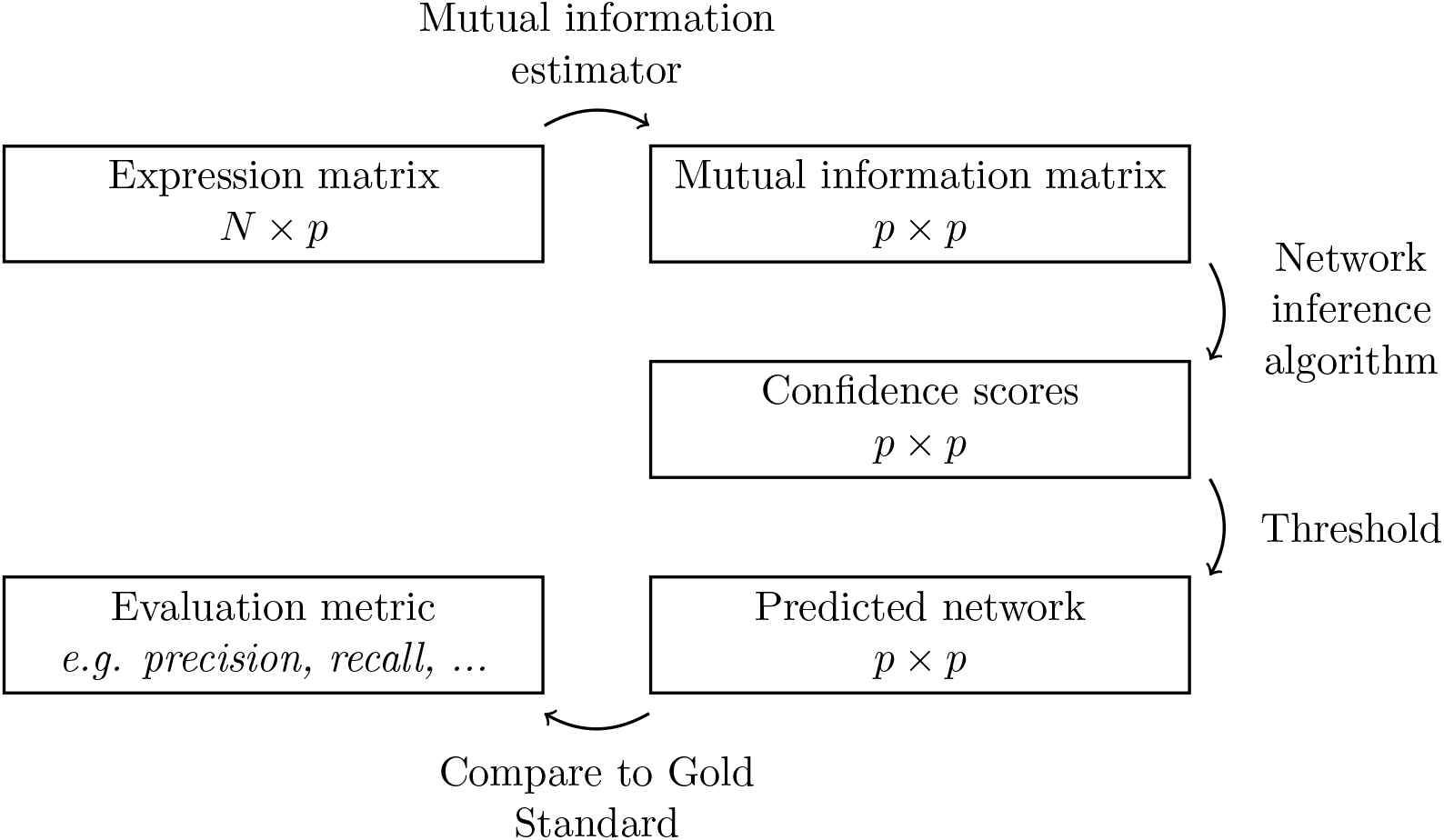
The network inference workflow for mutual information-based inference algorithms. Typically this process will be repeated using all possible thresholds, from which a precision-recall curve is plotted. The area under this curve is then used to assess the accuracy of the inferred network.

### 2.2 Mutual Information

Mutual information is an information-theoretic quantity that quantifies the dependence between two random variables. For two continuous random variables *x* and *y*, their mutual information is defined as

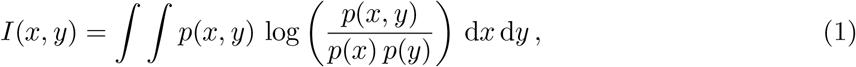

where *p*(*x, y*) is the joint probability distribution of *x* and *y* and *p*(*x*) and *p*(*y*) are the marginal probability distribution of *x* and *y* respectively. If *x* and *y* are discrete variables taking values in the sets *X* and *Y* respectively this definition becomes

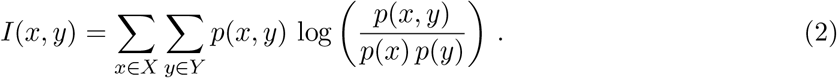

This definition can be equivalently expressed in terms of the (information theoretic) entropy as

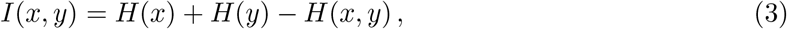

where *H*(*x*) and *H*(*y*) are the marginal entropies defined by

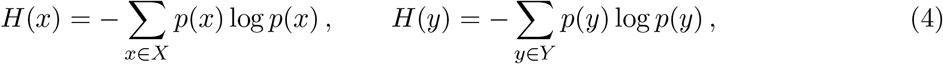

and *H*(*x, y*) is the joint entropy defined by

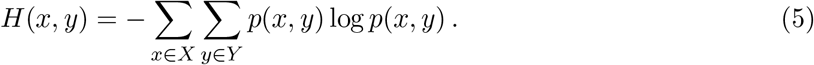

In practice, is common to use (3)-(5) to estimate *I*(*x, y*) if the mutual information estimator involves binning the expression data. In both (1) and (2) the base of the logarithm determines the units of *I*. Using log_2_ leads to a quantity measured in bits and the natural logarithm leads to a quantity measured in nats.

From (1), *I*(*x, y*) is symmetric and takes values in [0, *∞*). It is zero if and only if *x* and *y* are strictly independent, which is to say that *p*(*x, y*) = *p*(*x*) *p*(*y*). This enables the mutual information to capture non-linear dependencies between variables, unlike linear measures such as the Pearson correlation coefficient.

If *p*(*x, y*) is bivariate normal then the *I*(*x, y*) can be formulated exactly in terms of Pearson’s correlation coefficient

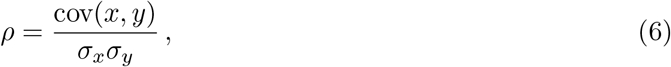

where cov is the covariance of *x* and *y* and *σ_x_, σ_y_* are the standard deviations of *x* and *y* respectively. In this case

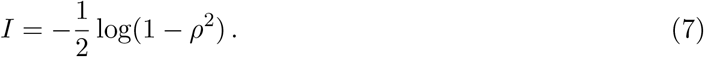

Another common approach uses Spearman’s correlation coefficient, where *x* and *y* are replaced by a ranking of their values.

### 2.3 Mutual information estimation

An accurate estimate of the mutual information requires accurate estimates of the distributions *p*(*x*), *p*(*y*) and *p*(*x, y*) from a finite (and often small) number of samples. Different mutual information estimators are essentially different approaches to estimating these distributions and each performs well in different scenarios. These different performance properties make different estimators suited to different tasks.

For example, many of the applications of mutual information come from machine learning, where mutual information can be used for feature selection [27] and independent component analysis [28]. Mutual information is also used in medical image analysis to align two images to the same coordinate system [29]. These two machine learning applications are concerned with finding independent random variables, and hence the mutual information estimator must be accurate near values close to zero. Medical image registration requires finding a coordinate system that maximises mutual information between images, and so the estimator must be accurate for large values. This is the regime which is important for network inference, where we are concerned with detecting strong dependencies between the expression of two genes.

One approach to estimating mutual information is to place the data into bins and use histograms to estimate the required distributions. This simple approach is the Maximum Likelihood estimator and is known to be negatively biased [30]. The Miller-Madow estimator [31] adds a correction to correct this bias, while the Shrinkage estimator [32] uses a uniform distribution to regularise the empirical distribution. Since network inference produces a ranked list of scores from mutual information estimates a constant bias added to each value should not affect the inferred network. The performance of these histogram-based estimators depend heavily on an appropriate choice of origin, the number of bins and bin locations.

The Chao-Shen estimator [33] was developed to estimate the diversity of biological species from small sample sizes using entropy. It is adapted to account for species that are not included in a specific sample, which corresponds to an empty bin. When using the Chao-Shen estimator for network inference, continuous expression data is binned using parameters that are decided by the user. Since the user controls the binning of expression data it is not clear how the Chao-Shen estimator’s design motivations are applicable for network inference, but it is included as it widely used and has a straightforward implementation.

Binning continuous data is a hard operation that is sensitive to noise. The B-spline, Kernel Density and *k*-nearest-neighbour estimators use smoothing methods to obtain more accurate estimates of the distributions. The B-spline estimator [8] is an adaptation of the Maximum Likelihood estimator that allows data points to be placed in multiple bins, thus smoothing the distribution. This procedure also reduces the dependence of the result on the choice of origin and the bin locations. The Kernel Density estimator [34] estimates the distributions using kernel functions rather than rectangular bins. Finally, the *k*-nearest-neighbour estimator [35] estimates the distribution using distances to nearest neighbours. Two additional estimators were included that use the Spearman and Pearson correlation with (7) to calculate the mutual information.

The parameters of the various estimators are shown in Table 2. Further information on the estimators and their parameters is given in Section S1.

**Table 2:** The parameters of the different mutual information estimators. See Section S1 for further details on the estimators and their parameters. The parameters of the Kernel Density estimator were not investigated.

| *Estimator(s)* | *Parameters* |
| --- | --- |
| Maximum Likelihood, Miller-Madow,<br>Chao-Shen, Shrinkage | Number of bins, binning method |
| B-spline | Spline order, number of bins |
| Kernel Density | Bandwidth, kernel (not investigated) |
| $k$-nearest-neighbour | $k$ |
| Spearman, Pearson correlation | n/a |

## 3 Results

### 3.1 Comparing mutual information estimators

A parameter study, which is discussed in Section 3.2, attempted to identify parameters that led to large AUPRC values across all the datasets. Separate parameters were identified for each inference algorithm. These are displayed in Table 3. As the best performing inference algorithm, CLR will be the focus of this discussion.

**Table 3:**
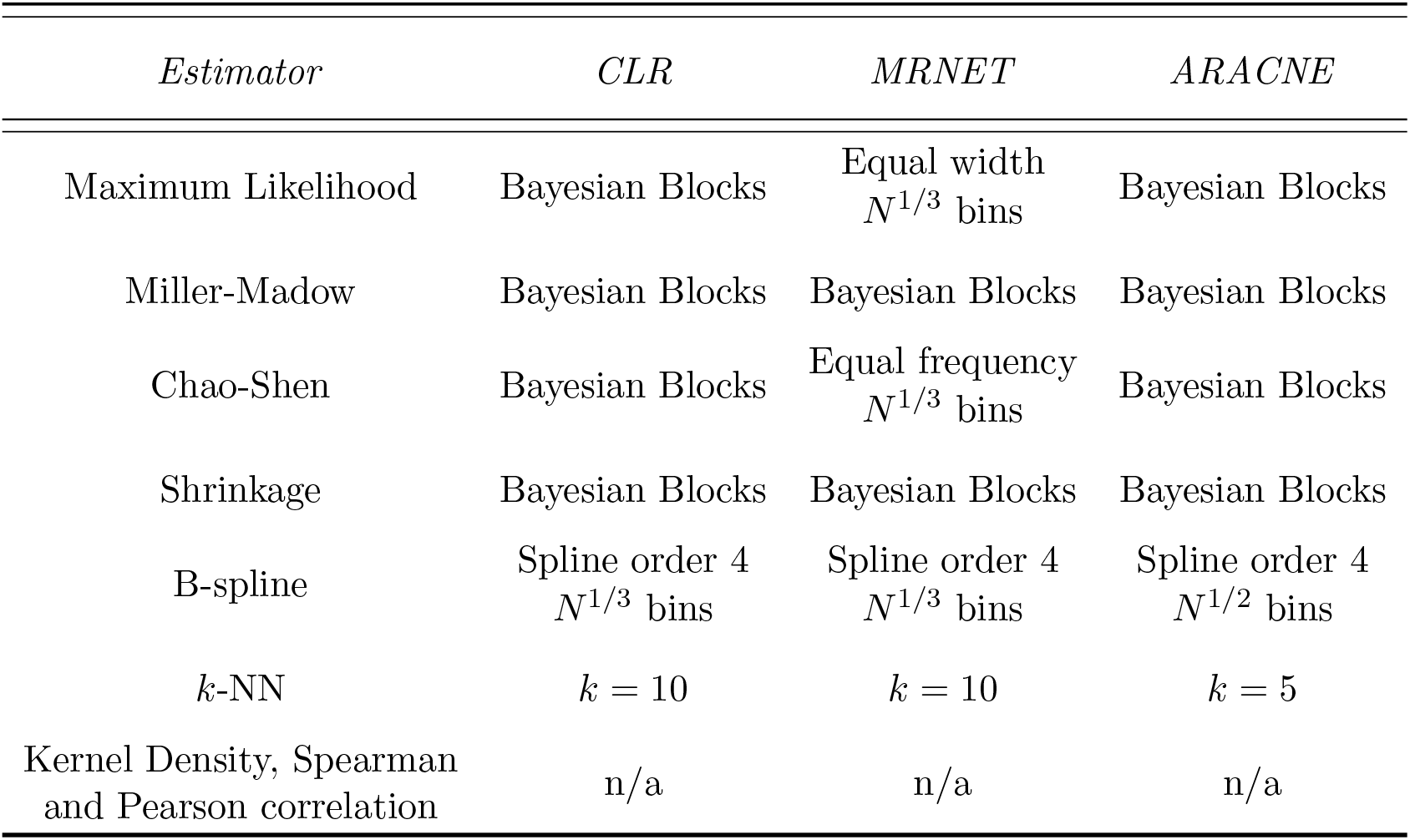
The parameters that maximise the AUPRC for each combination of inference algorithm and mutual information estimator. These parameters were chosen following the investigation whose results are presented in Section S5. How these parameters were chosen is discussed in Section 3.2. The parameters of the Kernel Density estimator were not investigated and the only the values suggested by the authors of [34] were used. The Spearman and Pearson correlation estimators do not have parameters.

#### 3.1.1 *in silico* dataset

The AUPRC values for the *in silico* dataset are shown in Figure 2. CLR gives the largest AUPRC values while ARACNE gives the lowest. This is the case for all three datasets. Furthermore, the spline estimator gives the largest AUPRC for both CLR and MRNET. For MRNET the benefit of using the B-spline estimator means that it performs better than CLR with many of the mutual information estimators. This is also true of MRNET with the Kernel Density estimator, but the AUPRC is lower than using MRNET with B-spline.

**Figure 2:**
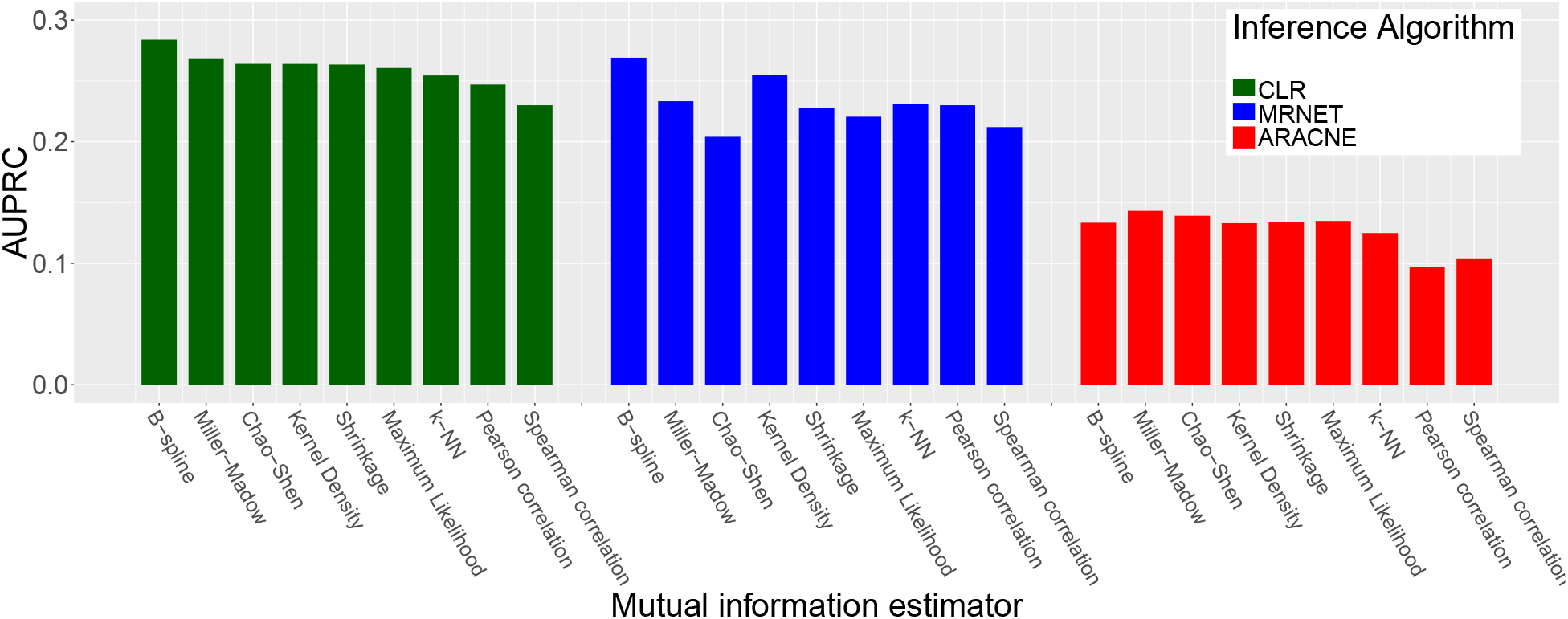
The resulting area under precision-recall curve (AUPRC) when using the different mutual information estimators with each of the inference algorithms on the *in silico* dataset. The parameters for each estimator are those which were found to maximise the AUPRC following the investigation described in Section S5. The results of this investigation can be found in Section S5.

#### 3.1.2 *E. coli* dataset

The AUPRC for the different mutual information estimators is shown in Figure 3. The AUPRC values are lower for the *E. coli* dataset than for the *in silico* dataset, which reflects the added difficulty of inferring a network and obtaining an accurate gold standard for real biological systems (see Section 4.1 for discussion on this topic). For this dataset there is not an estimator that performs significantly better than the others. However, the Kernel Density estimator performs significantly worse than the other estimators for all three inference algorithms, and MRNET in particular.

**Figure 3:**
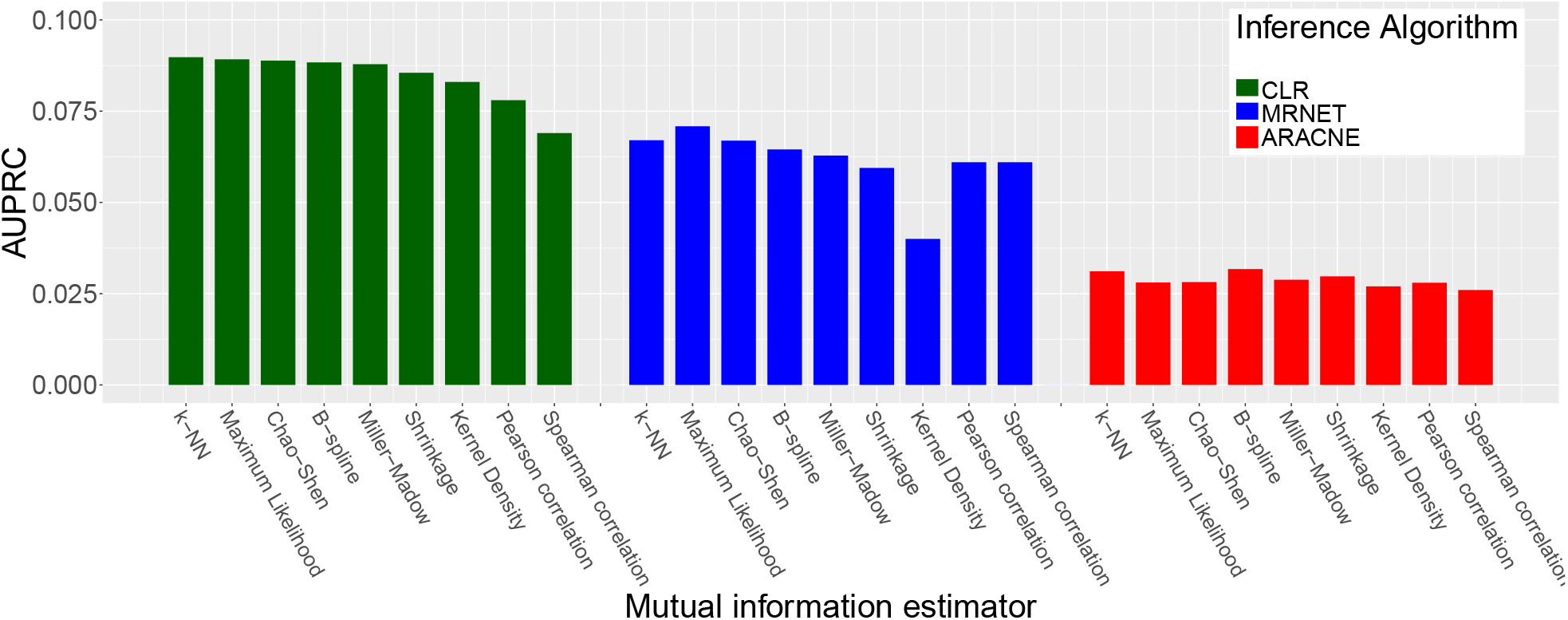
The resulting area under precision-recall curve (AUPRC) when using the different mutual information estimators with each of the inference algorithms on the *E. coli* dataset. The parameters for each estimator are those which were found to maximise the AUPRC following the investigation described in Section S5. The results of this investigation can be found in Section S5.

#### 3.1.3 *S. cerevisiae* dataset

The AUPRC for the different mutual information estimators is shown in Figure 4. For this dataset the variation in AUPRC between the estimators is negligible. The parameter study also found negligible variation in AUPRC with the parameters of all the estimators whose parameters were investigated (not the Kernel Density estimator). Possible explanations are discussed in Section 4.2.

**Figure 4:**
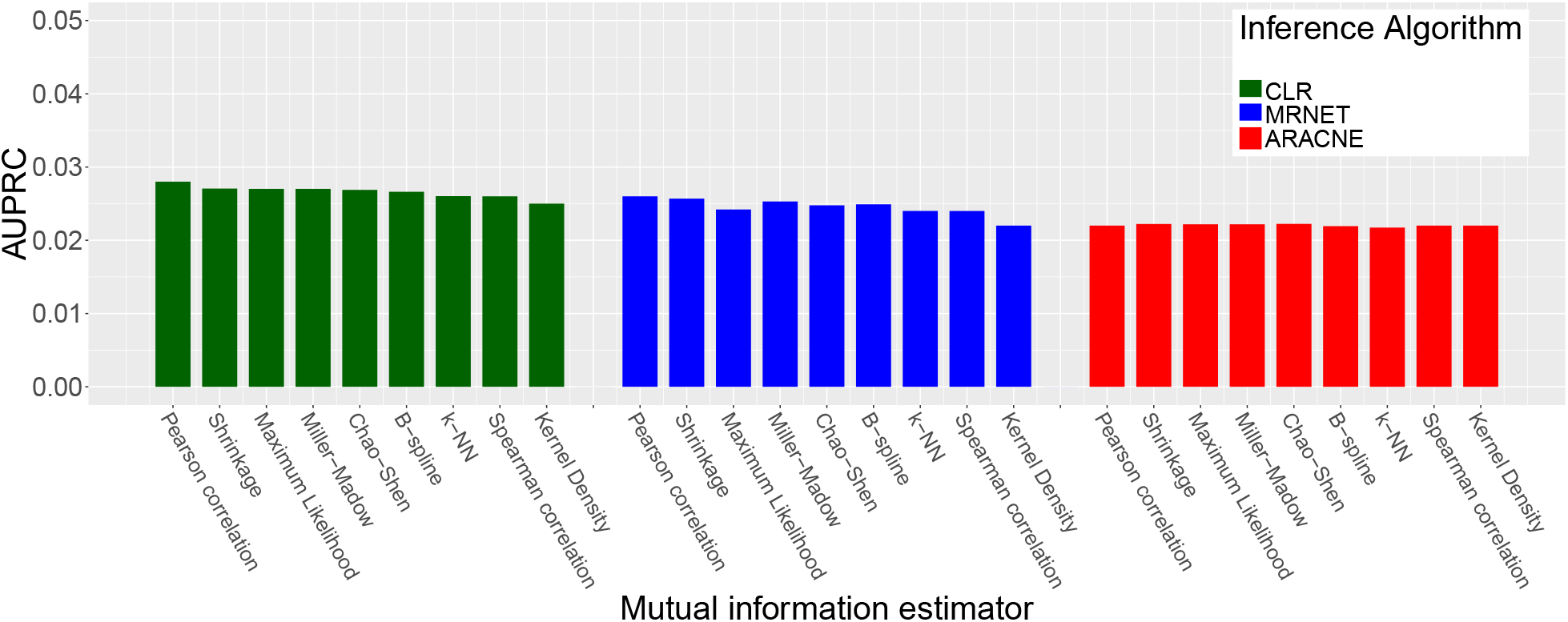
The resulting area under precision-recall curve (AUPRC) when using the different mutual information estimators with each of the inference algorithms on the *S. cerevisiae* dataset. The parameters for each estimator are those which were found to maximise the AUPRC following the investigation described in Section S5. The results of this investigation can be found in Section S5.

### 3.2 Mutual information estimator parameters

Part of this study examined estimator parameters with the aim of identifying optimal parameters that maximise the AUPRC. The full results can be found in Section S5.

It was found in [7] that the optimal parameters of the Maximum Likelihood, Miller-Madow and Shrinkage estimators are different for CLR, MRNET and ARACNE. Therefore it is expected that the optimal parameters will be different for the three inference algorithms. If possible, parameters were chosen that gave large AUPRC values on all three datasets. If parameters led to good performance on just the *in silico* and *E. coli* datasets then they were chosen. The *E. coli* dataset was given preference as its AUPRC results were found to vary more with parameters than the *S. cerevisiae* results.

The parameters that maximise the AUPRC when using CLR are shown in Table 4. CLR was included here as it was the best-performing inference algorithm on all three datasets (see Section 3.1), but equivalent results for MRNET and ARACNE are available in Section S6. Table 4 shows that there is more consistency within datasets than within estimators.

**Table 4:**
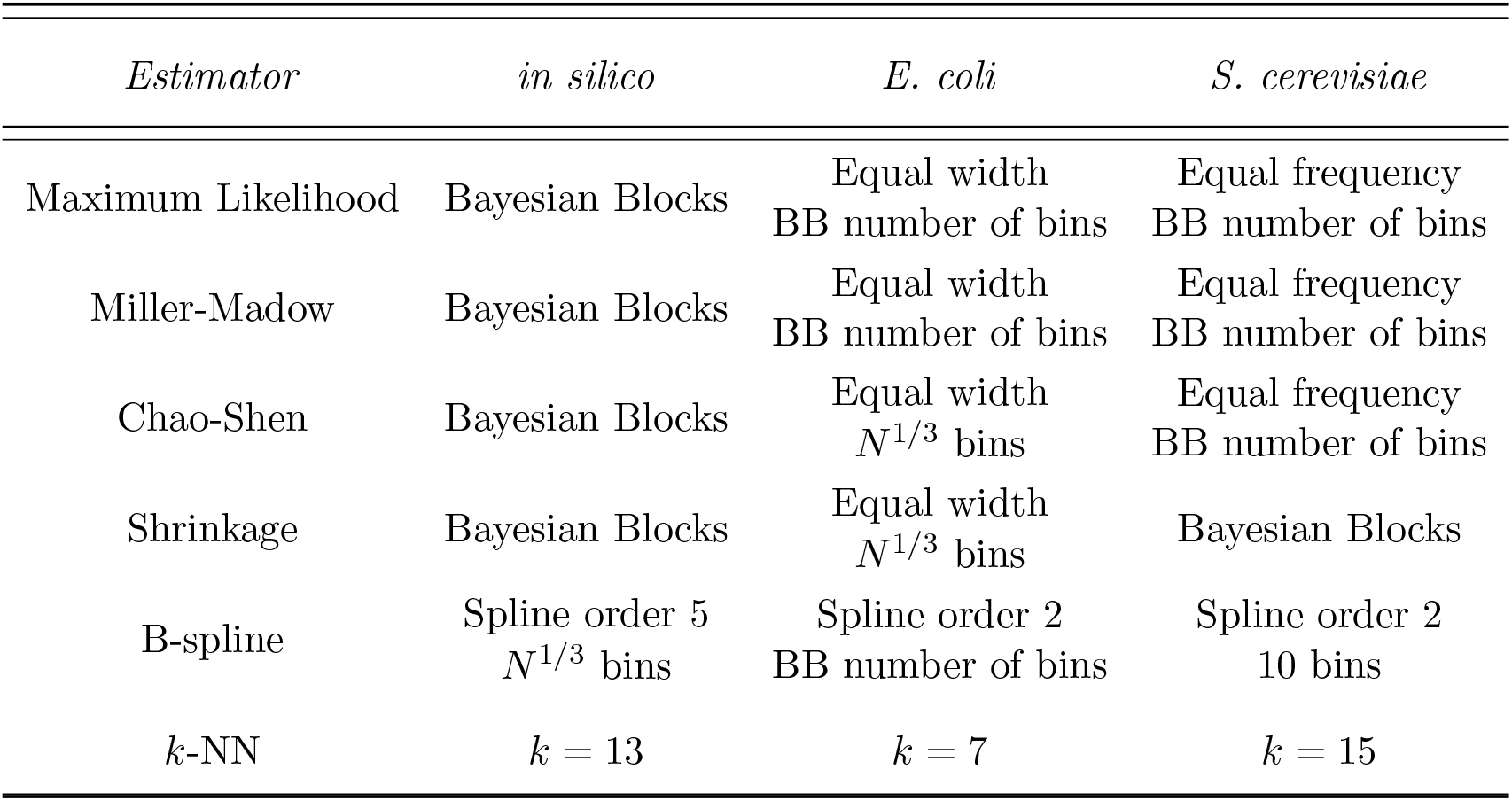
The mutual information estimator parameters that maximise the AUPRC when using the CLR network inference algorithm. The Spearman and Pearson correlation estimators do not have parameters, while the parameters of the Kernel Density estimator were not investigated. The parameters of each estimator are shown in Table 2 and described in detail in Section S1.

#### 3.2.1 Number of bins for histogram-based estimators

This parameter study identified the number of bins as the key parameter of the Maximum Likelihood, Miller-Madow, Chao-Shen and Shrinkage estimators. Common choices are 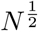 bins or 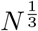 bins, which lead to the same number of bins for all genes. Another option is the Freedman-Diaconis rule [36], which uses bins with size 2. 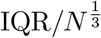, where IQR is the inter-quartile range of the expression profile of a gene [36]. Note that this leads to different numbers of bins for each gene. The Freedman-Diaconis rule resulted in low AUPRC values and its results are not included here. See the Section S5 for the full results. Since the Bayesian Blocks algorithm chooses the number of bins automatically, another option is to use the number of bins chosen by Bayesian Blocks but with equal frequency or equal width bins.

Both Bayesian Blocks and Freedman-Diaconis bins choose the number of bins for each gene, resulting in a distribution. These distributions are shown for each dataset in Figure 5. The Freedman-Diaconis rule produces results that are approximately centred at *N* ^1*/*2^. For Bayesian Blocks the resulting distribution is closer to *N* ^1*/*3^ and also has a smaller variance than the corresponding distribution for the Freedman-Diaconis rule.

**Figure 5:**
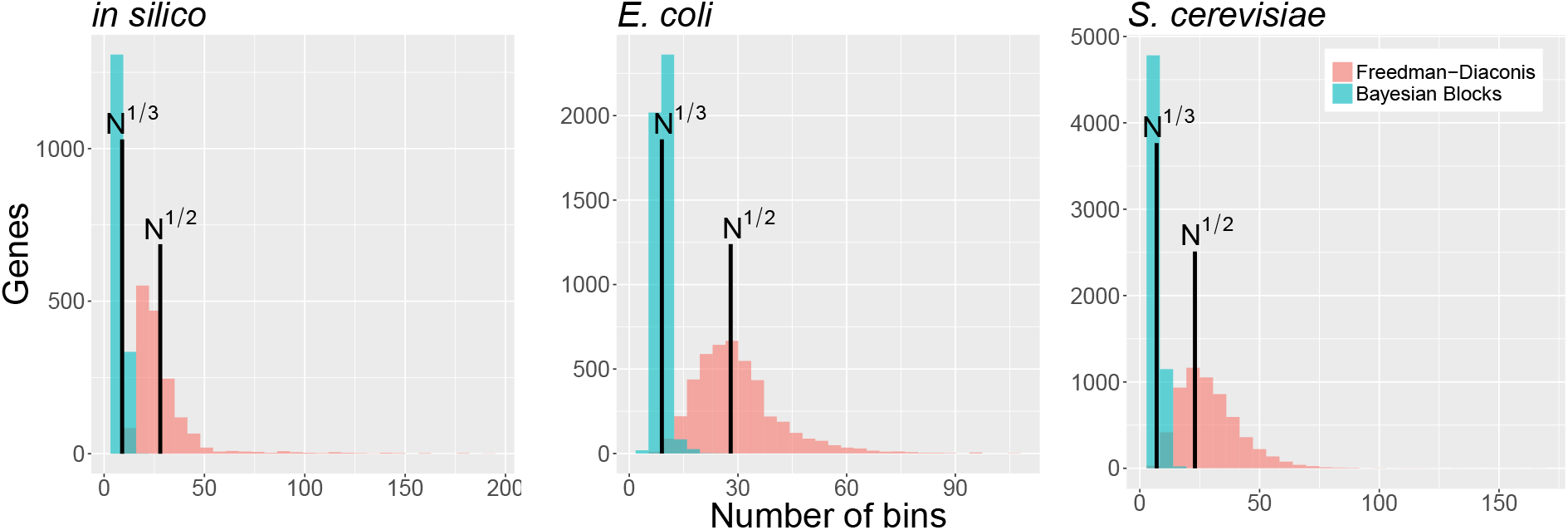
The resulting number of bins for each gene in the three datasets. The Freedman-Diaconis rule and Bayesian Blocks choose different numbers of bins for each gene, while the using *N* ^1*/*2^ or *N* ^1*/*3^ bins gives the same number for all genes. 16 genes with large numbers of Freedman-Diaconis bins have been excluded from the *in silico* plot for aesthetic reasons.

It is expected that the AUPRC from *N* ^1*/*3^ or the number of bins from Bayesian Blocks should give a similar AUPRC as they result in a similar number of bins (see Figure 5). This is observed in Figures 6 and 7, which show that these two bin number rules give a larger AUPRC than *N* ^1*/*2^ bins for the *in silico* and *E. coli* datasets. The relative increase in AUPRC is smaller for the *E. coli* dataset than *in silico*. These results also show that the AUPRC is more robust to the Binning Method than the number of bins. For the *S. cerevisiae* dataset, whose results are shown in Figure 8, there is not a large variation in AUPRC between the different bin number rules. The lack of variation in AUPRC between estimators and parameters for the *S. cerevisiae* dataset will be discussed in Section 4.2.

**Figure 6:**
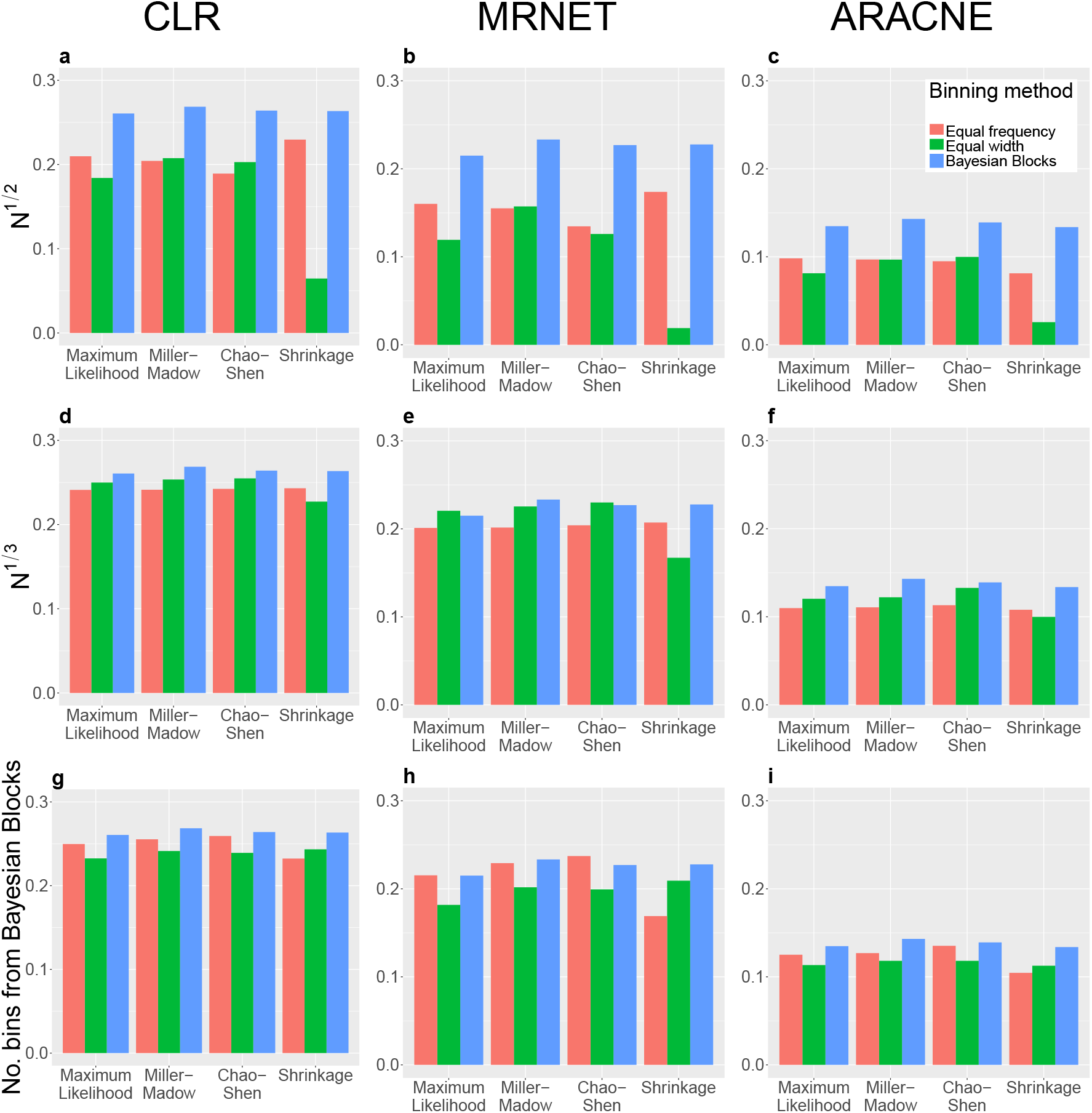
The AUPRC for the *in silico* dataset with the Maximum Likelihood, Miller-Madow, Chao-Shen and Shrinkage estimators. An individual plot shows the AUPRC when using a single inference algorithm and a specific number of bins with the four estimators. Each grouping of bars represents a single estimator for the three binning methods. Each column shows results for a single inference algorithm and each row shows results for a single number of bins. Note that when using the Bayesian Blocks binning method the number of bins is chosen automatically, hence the AUPRC values within columns are the same for the same MI estimator, but are included in all plots for ease of comparison. The Freedman-Diaconis rule has been removed from this plot as its AUPRC were signi_cantly lower than the other number of bins but are included in Figure S2.

**Figure 7:**
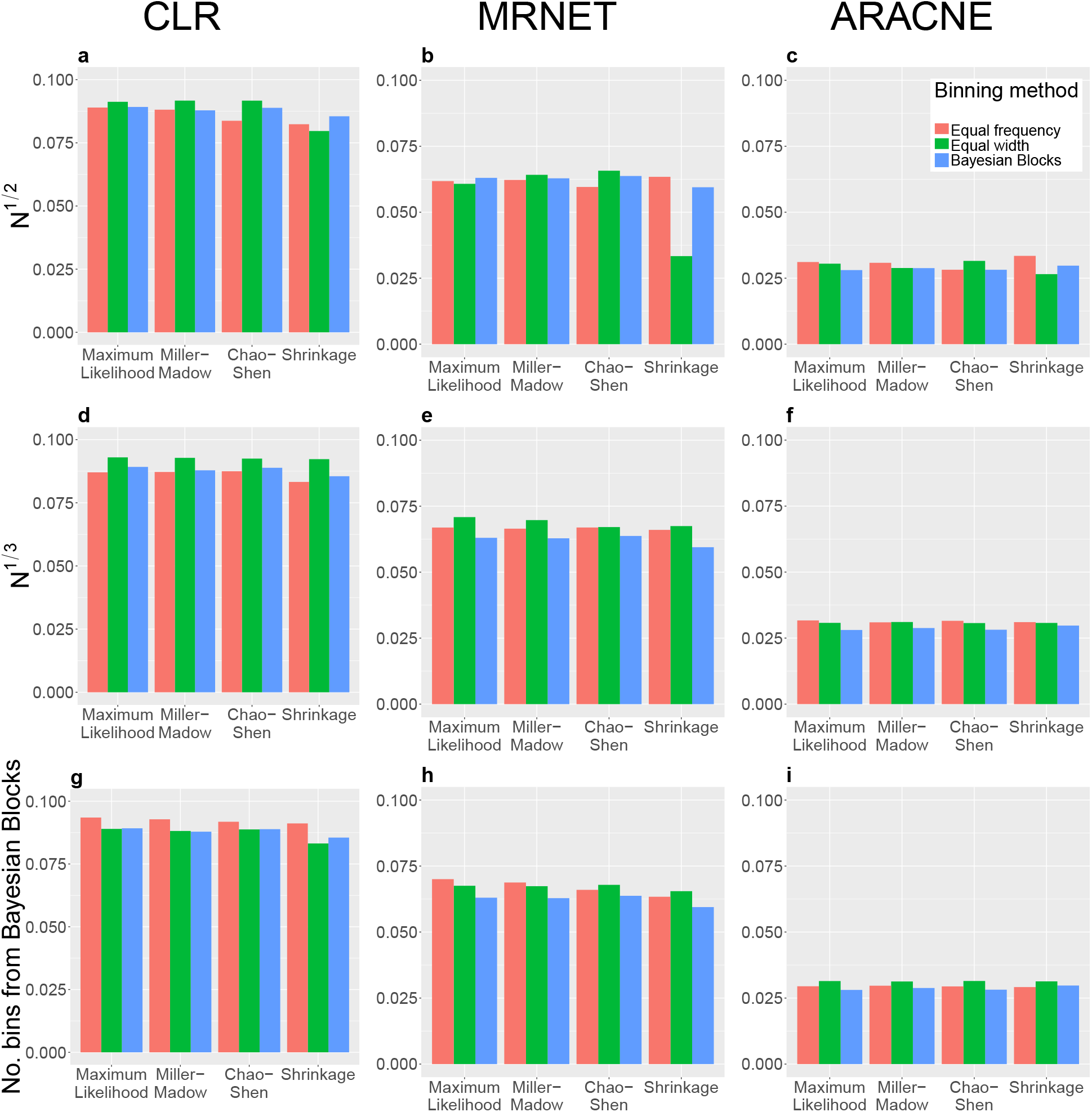
The AUPRC for the *E. coli* dataset with the Maximum Likelihood, Miller-Madow, Chao-Shen and Shrinkage estimators. An individual plot shows the AUPRC when using a single inference algorithm and a specific number of bins with the four estimators. Each grouping of bars represents a single estimator for the three binning methods. Each column shows results for a single inference algorithm and each row shows results for a single number of bins. Note that when using the Bayesian Blocks binning method the number of bins is chosen automatically, hence the AUPRC values within columns are the same for the same MI estimator, but are included in all plots for ease of comparison. The Freedman-Diaconis rule has been removed from this plot as its AUPRC were signi_cantly lower than the other number of bins but are included in Figure S5.

**Figure 8:**
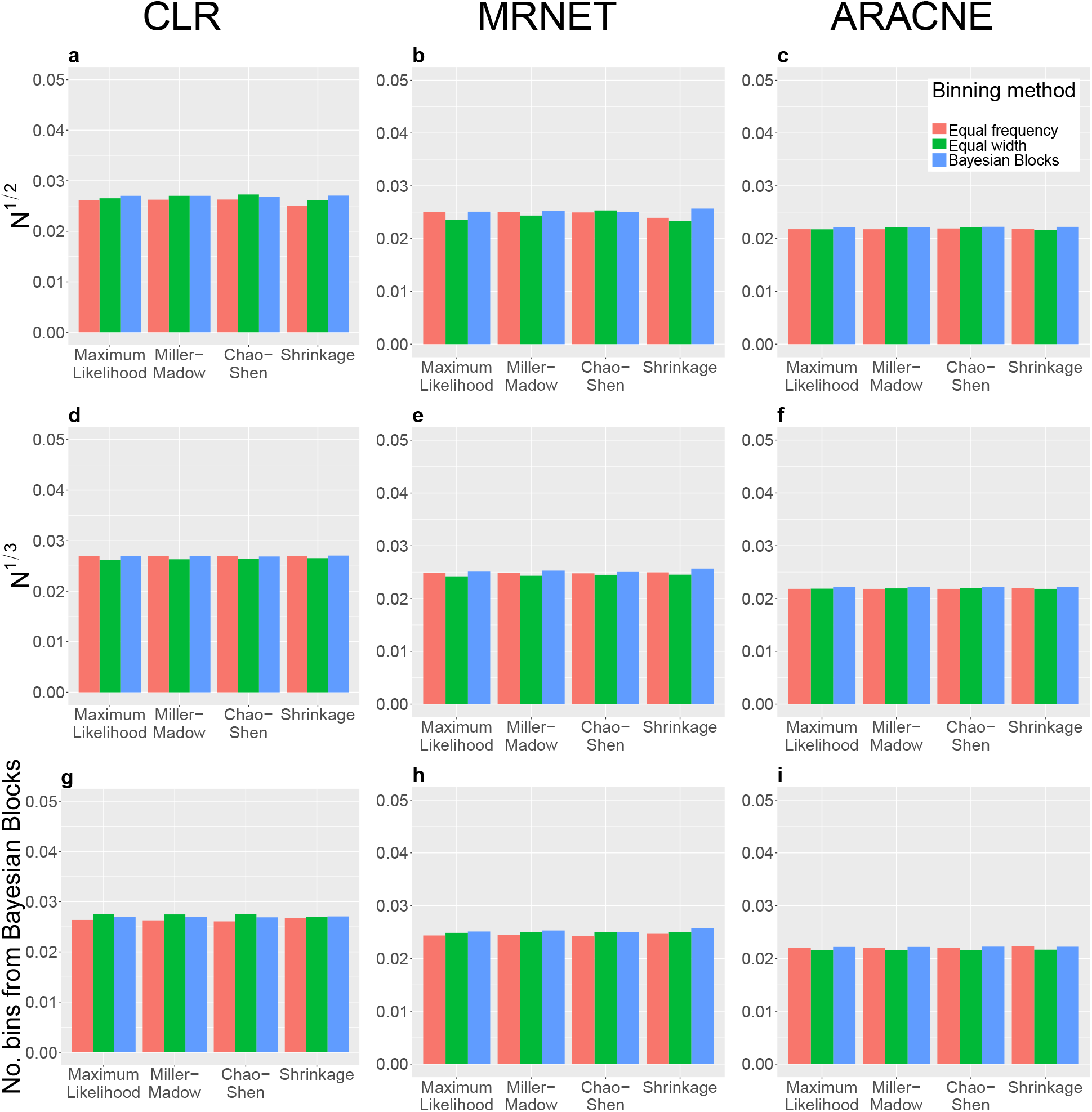
The AUPRC for the *S. cerevisiae* dataset with the Maximum Likelihood, Miller-Madow, Chao-Shen and Shrinkage estimators. An individual plot shows the AUPRC when using a single inference algorithm and a specific number of bins with the four estimators. Each grouping of bars represents a single estimator for the three binning methods. Each column shows results for a single inference algorithm and each row shows results for a single number of bins. Note that when using the Bayesian Blocks binning method the number of bins is chosen automatically, hence the AUPRC values within columns are the same for the same MI estimator, but are included in all plots for ease of comparison. The Freedman-Diaconis rule has been removed from this plot as its AUPRC were significantly lower than the other number of bins but are included in Figure S8.

These findings indicate that using the number of bins found by Bayesian Blocks improves the accuracy of inferred networks, even if the bin locations found by Bayesian Blocks are not used. A simpler alternative is to use *N* ^1*/*3^ bins, which is typically close to the number of bins used by Bayesian Blocks.

### 3.3 Computation times

The computation time of each estimator is clearly an important practical consideration. The number of gene pairs is *𝒪*(*p*^2^) for *p* genes and linear in the number of samples. Table 5 shows the computation time of several implementations of the mutual information estimators used in this study. In producing this work several estimators were implemented using parallel C++ with an R interface and are available in the R package *fastGeneMI*. The package can be downloaded at https://bitbucket.org/Jonathan-Ish-Horowicz/fastgenemi/. Table 5 also includes the computation times when using alternative implementations that compute mutual information estimates from a matrix of continuous expression values. Another R package, “synRNASeqNet,” enables the calculation of mutual information estimates from binned data only and so was not included. Note that using the Bayesian Blocks binning method with the datasets from this study requires additional computation time of order 100 seconds.

**Table 5:** Time to compute the mutual information between the expression of all gene pairs for the *E. coli* dataset (4,297 genes, 805 samples) with different implementations of the estimators used in this study. Results are given in seconds, with the lowest time in bold text. In brackets is the computation time relative to the fastest implementation for that estimator. Computations were performed by 16 2.6 GHz Intel Sandy Bridge cores. If an implementation allows parallel computation then all 16 cores were used. n/a indicates that a package does not provide an implementation for that estimator.

| *Estimator* | *fastGeneMI* | *minet* [37] | *DepEst* [38] | *parmigene* [39] |
| --- | --- | --- | --- | --- |
| Maximum Likelihood | **6.6 (1.0)** | 23132.0 (3,120.1) | n/a | n/a |
| Miller-Madow | **6.6 (1.0)** | 23,180.3 (3,509.0) | 28.4 (4.3) | n/a |
| Chao-Shen | **10.7 (1.0)** | n/a | 28.513 (2.7) | n/a |
| Shrinkage | **6.0 (1.0)** | 23,271.0 (3,885.6) | n/a | n/a |
| B-Spline | **101.4 (1.0)** | n/a | 178.1 (1.8) | n/a |
| Kernel Density | n/a | n/a | **75,188.0 (1.0)** | n/a |
| *k*-NN | n/a | n/a | >86,400.0 (>211.4) | **408.8 (1.0)** |

The histogram-based estimators have the shortest computation times, followed by the B-spline estimator. The computation time of the B-spline estimator is an order of magnitude larger. The Kernel Density and *k*-nearest-neighbour estimators have computation times two orders of magnitude larger than the histogram-based methods.

## 4 Discussion

### CLR was the best performing inference algorithm

The results in Section 3.1 provide further evidence that CLR is the state of the art mutual-information based inference algorithm [1, 7]. Furthermore, it has been shown here to be robust to both the choice of estimator and the estimator parameters. This was evident across all the estimators whose parameters were investigated (see Section S5). MRNET is able to match the performance of CLR, but this requires specific choices of estimator and parameters.

### The B-spline estimator was the best performing mutual information estimator

We recommend using the B-spline estimator, supporting the findings of [9, 22]. The AUPRC from the B-spline was among the largest for the *in silico* and *E. coli* datasets for each inference algorithm used in this study. The B-spline estimator performed less well on the *S. cerevisiae* dataset, however the variation in AUPRC for this dataset was smaller than the other two. The B-spline computation time is also significantly lower than for other smoothing estimators (Kernel Density and *k*-nearest-neighbour). The computation time is longer than the other histogram-based estimators, but these lead to lower AUPRC values. This may be because the binning of data is a hard operation and is thus extremely sensitive to noise and the locations of the bins. The B-spline estimator reduces this sensitivity by placing samples in multiple bins. The B-spline estimator was also found to be robust to changes in its parameters (see Figures S3, S6 and S9).

### When using an estimator that bins expression data, use *N* ^1*/*3^ bins for *N* samples

These results indicate that when using a mutual information estimator that bins the data, using *N* ^1*/*3^ bins is the optimal choice. This contradicts a previous finding by Kurt et al. who reported that *N* ^1*/*2^ bins led to the most accurate inferred network [22]. These differing results may be because that study used a F-score as the performance metric. We believe that AUPRC is a more appropriate metric for network inference and therefore that *N* ^1*/*3^ is a superior choice of bin number. As discussed in Section 2.1, CLR, MRNET and ARACNE produce a list of edges that are ranked by their corresponding scores. To produce a single network from this ranked list a threshold is chosen, and any edge with a score greater than the threshold is designated as an edge. The F-score can only be evaluated for a single network and is therefore highly dependent on the choice of threshold. The AUPRC is computed using all possible thresholds by definition and so removes this dependency.

### Bayesian Blocks did not improve the accuracy of the inferred networks on the real datasets

For the *in silico* dataset, using Bayesian Blocks bins led to larger AUPRC values than equal frequency or equal width bins. However, this was not the case for either of the real dataset. A possible explanation for this difference lies in the different processing of the synthetic and real expression data. The *in silico* expression data are raw values as simulated by GeneNetWeaver, while the real expression data have undergone microarray normalisation followed by a log transform. The expression of a single gene of the real datasets is therefore more normal than a gene from the synthetic dataset, which has a large number of zeros. This can be seen by comparing Figures 9 and 10. Figure 9 shows a concentration of values at zero. Bayesian Blocks represents these values using a narrow bin with a high frequency density, but equal frequency and equal width bins do not represent this feature of the data well. Real expression data (Figure 10) does not have this feature, and the three binning methods give qualitatively similar bins. For more information on how the expression data was prepared see the Supplementary Material of [1].

**Figure 9:**
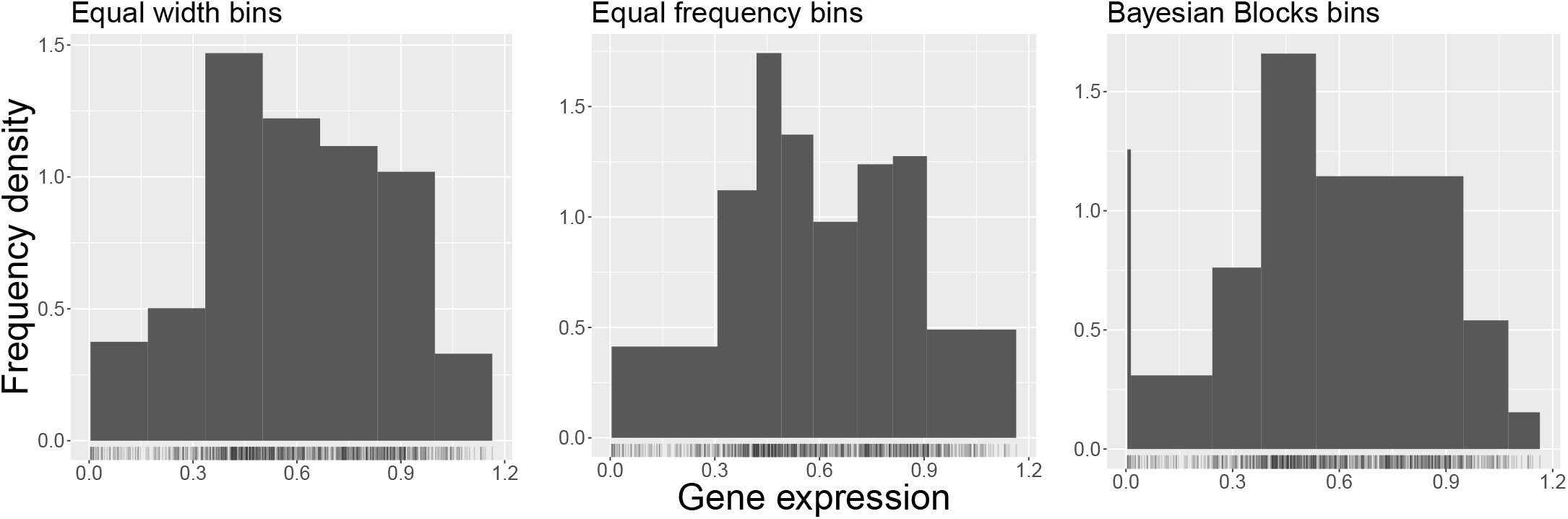
Histograms of the expression of the gadE gene (index 126) of the *in silico* dataset using the three binning methods. For the equal width and equal frequency methods the number of bins were chosen to match those used in the Bayesian Blocks case. For the unprocessed synthetic data there are are large number of zeros which are best represented by the Bayesian Blocks bins.

**Figure 10:**
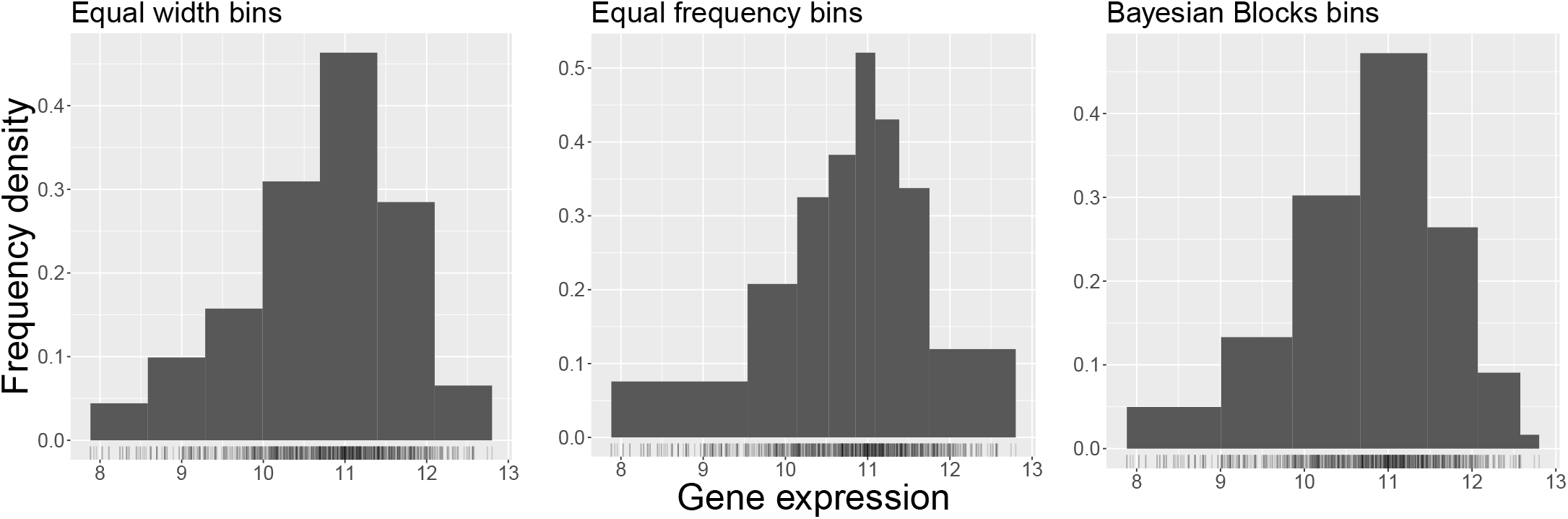
Histograms of the expression of the secE gene (index 1263) of the *E. coli* dataset using the three binning methods. For the equal width and equal frequency methods the number of bins were chosen to match those used in the Bayesian Blocks case. The three binning methods give more similar results for the processed real data than for the synthetic data (Figure 9).

For all the datasets the number of bins found by Bayesian Blocks was close to *N* ^1*/*3^ Furthermore, since the recommended inference algorithm-mutual information estimator combination (CLR-B-spline) is robust to the number of bins, we recommend using *N* ^1*/*3^ bins.

### 4.1 The relationship between synthetic and real expression data

These results raise questions about the relationship between synthetic and real expression data. Synthetic expression data generators are valuable and widely used tools for benchmarking network inference algorithms. However, the AUPRC values on the *E. coli* and *S. cerevisiae* datasets were significantly lower than on the *in silico* dataset. Furthermore, the AUPRC values for the *in silico* exhibited greater sensitivity to both the choice of estimator and estimator parameters.

There are many causes of the decrease in AUPRC when moving from a synthetic to a real datasets. Measuring expression using DNA microarrays requires a set of complex steps, each of which introduce noise. Each of these steps involves several choices that are also known to impact on the measured expression values, such as the choice of normalisation procedure and the sample procedure [40]. Although synthetic datasets allow the user to control additive noise it is not clear what is an appropriate noise amplitude to choose. It is also unlikely that additive noise is a good approximation of the noise resulting from a series of complex experimental techniques.

An additional source of the discrepancy in AURPC between the real and synthetic data concerns the gold standard network. While the synthetic gold standard is known exactly, the real gold standard networks contain many false negatives. For example, the DREAM5 Network Inference Challenge identified 53 *E. coli* potentially novel interactions using a consensus of predictions by arange of inference algorithms. These were edges that were not present in the gold standard networks but were confidently predicted as edges by a number of inference algorithms. Of these interactions, 23 (43%) were subsequently verified experimentally [1]. Furthermore, since the networks used by GeneNetWeaver are sub-networks of real biological networks they are intrinsically biased against undiscovered regulatory links. That is to say, their potential to aid in the discovery of novel regulatory interactions is limited since their network topologies are based on incomplete biological knowledge. These comments do not aim to denigrate their use in the development of novel inference algorithms, simply to note that there remains a significant discrepancy between the inference of synthetic and real transcriptional regulatory networks.

### 4.2 Comparing the results of the *E. coli* and *S. cerevisiae* datasets

A similar trend was observed between the two real datasets. The *E. coli* dataset AUPRC statistics were larger and more dependent on the estimator and its parameters than the *S. cerevisiae* dataset results. One cause may be that this dataset contains 563 samples while the *E. coli* dataset contain 805 samples. The lower number of samples makes it more difficult to estimate the probability distributions required for an estimate of the mutual information, and so it may be that all of the estimators are producing inaccurate mutual information estimates.

There may also be a biological explanation for the discrepancy in results between the two real datasets. *E. coli* is a prokaryote and is therefore less complex than the eukaryotic *S. cerevisiae*. The added complexity of the *S. cerevisiae* transcriptional regulatory network is most likely a primary cause of the lower AURPC results. The lack of variation of AURPC for the *S. cerevisiae* dataset could suggest that mutual information-based inference algorithms have difficulty inferring its transcriptional regulatory network. However, the AUPRC values found in [1] are essentially constant across several other types of inference algorithm. This suggests either that the currently available approaches are fundamentally flawed, or that the *S. cerevisiae* gold standard is significantly more incomplete than that of *E. coli*. Specifically, the integration of heterogeneous data sources may be required for an accurate reconstruction of the *S. cerevisiae* transcriptional regulatory network [41]. Another possible explanation is that a larger part of the transcriptional regulation in eukaryotes is performed by complexes of transcription factors and other proteins, which are impossible to identify using only pairwise interactions.

## Supplementary Material

### S1 Descriptions of Estimators

This section consists of a detailed description of the MI estimators used in this study. As mentioned above, to estimate *I*(*x, y*) requires estimates of the marginal distributions *p*(*x*), *p*(*y*) and the joint distribution *p*(*x, y*). Estimates of these distributions will be denoted with as *p̂*

The Maximum Likelihood, Miller-Madow, Chao-Shen, Shrinkage and B-spline estimators were implemented in parallel C++ with an R interface. These implementations are available in the R package *fastGeneMI*. The package can be downloaded from https://bitbucket.org/Jonathan-Ish-Horowicz/fastgenemi/. The *depEst* implementation [38] of the Kernel Density estimator was used with default parameters throughout. The *k*-nearest-neighbour estimator implementation from the *parmigene* R package was used [39].

#### S1.1 Maximum Likelihood Estimator

The first histogram-based method is the Maximum Likelihood estimator, which discretises the continuous expression data into bins then approximates the marginal probability distributions using

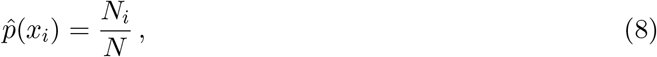

where *x_i_* are the range of values covered by the *i*-th bin, *N_i_* is the number of samples that fall into bin *i* and *N* = Σ_*i*_*N_i_* is the total number of samples. Similarly, the estimate of the joint distribution is

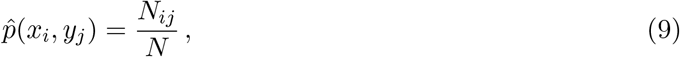

where *y_j_* is the range of the *j*-th bin in *y*, *N_ij_* are the number of samples in the *i*-th bin for *x* and the *j*-th bin for *y* and *N* = Σ_*i*_ Σ*_j_ N_ij_*. The Maximum Likelihood entropy estimates are then given by

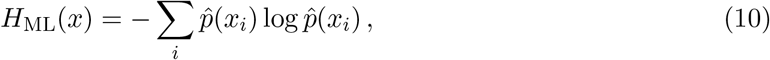

and

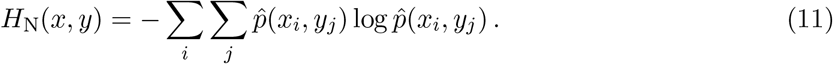

A mutual information estimate is then calculated using (3). (8) and (9) are often called “empirical distributions. The Maximum Likelihood estimator is known to be negatively biased, especially for small sample sizes [30]. It is also known as the “Empirical,” “Plug-in” or “Naive” estimator.

#### S1.2 Miller-Madow Estimator

Introduced in [31] by Miller, this estimator introduces an additional term to the entropy estimates (10) and (11) to correct for the negative bias of the Maximum Likelihood estimator. The entropy estimates are

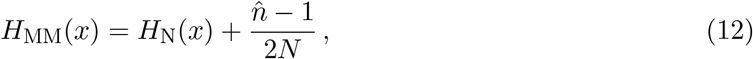

and

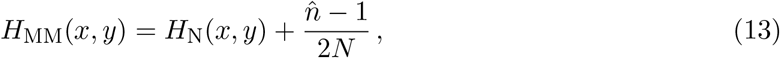

where *n̂* is the number of non-empty bins. A mutual information estimate is then found using (3).

#### S1.3 Chao-Shen Estimator

This estimator, proposed in [33] by Chao and Shen, was developed for the estimation of the diversity of biological species via their entropy. In this context each bin corresponds to a species. This entropy estimator attempts to correct for the fact that some species may not be included in a single set of samples from a population. This is equivalent to a bin being empty.

The first component is a Horvitz-Thompson estimator, which attempts to correct for varying proportions of observations within strata in a stratified sample of a target population [42]. If *N* samples have been drawn from a population then the probability that bin *i* has been included is 1 *−* (1 *− p*̂(*x_i_*))*^N^*. This probability is inverse weighted for each bin when calculating the entropy.

The second component of this estimator is a Good-Turing correction, which attempts to account for previously empty bins [43]. An approximate form of the correction is

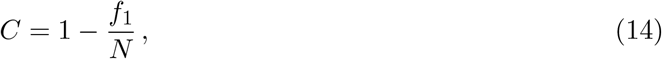

where *f*_1_ is the number of bins with a single count. Here we are assuming that the number of empty bins is the fraction of the bins with a single count. This correction is multiplied by all the estimates of the distributions in (4) and (5).

Together, these give the Chao-Shen entropy estimators

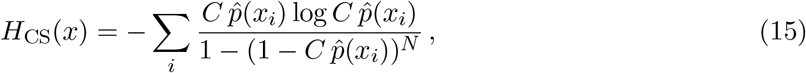

and

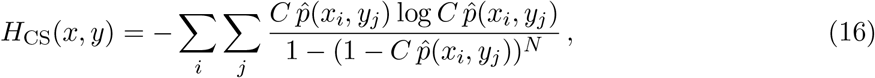

from which we can compute a mutual information estimate using (3).

For continuous gene expression data we choose the number and width of the bins, meaning that we control the number of empty bins. Therefore the motivation of the Chao-Shen estimator, which was developed for discrete data, is not directly applicable in the context of network inference.

#### S1.4 Shrinkage Estimator

This estimator, by Hausser and Strimmer [32], attempts to prevent overfitting using a convex sum of the empirical estimate of the distribution and a “target” distribution. For marginals, this “shrinkage” distribution is given by

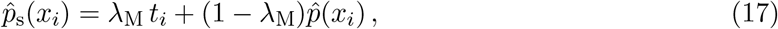

where *t_i_* is the target distribution and λ_M_ *∈* [0, 1] controls the relative weight of the two distributions. The target distribution is chosen to be the uniform distribution, 1*/*number of bins, which maximises the entropy. λ_M_ is given by

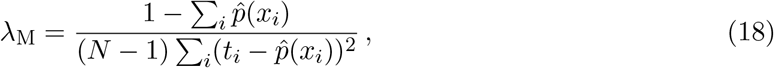

and is truncated to fall in [0,1]. The joint “shrinkage” distribution is

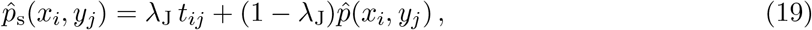

Where

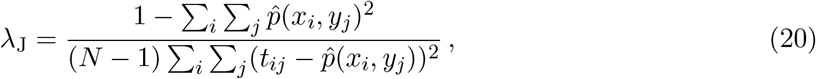

where the target distribution is once again 1*/*number of bins and λ_J_ *∈* [0, 1]. The shrinkage entropy estimates are

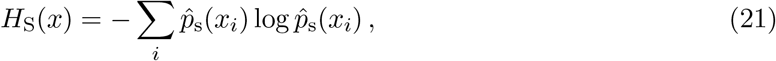

and

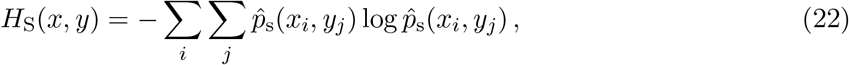

from which a mutual information estimate can be computed using (3).

#### S1.5 B-Spline Estimator

This method, proposed by Daub et al. in [8], is a modification of the Maximum Likelihood estimator that places samples in multiple bins, thus smoothing the distribution. This smoothing is achieved using B-splines.

A B-spline is a piecewise polynomial function, with each polynomial being a linear combination of so-called “basis functions.” In a B-spline of order *k* these basis functions are polynomials with degree *k −* 1. The piecewise polynomials are joined at the elements of a *knot vector t*, which has *k* + *m* elements when smoothing *m* bins using a B-spline of order *k*. The elements of the knot vector are given by

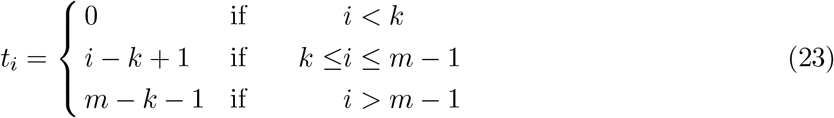

The knot vector is non-decreasing and fully determines the B-spline basis functions. For a B-spline of order *k* the basis functions are defined between *k* consecutive knots. The basis functions are determined recursively from *k* = 0 until the desired spline order is reached. For *z ∈* [0, *m−k* +1], the range of the knot vector, the *m* basis functions (one basis function per bin) are given by the Cox-de Boor recursion formula [44]:

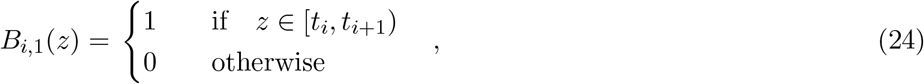

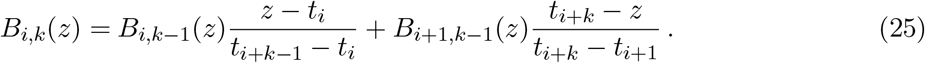

For *k* = 1 the basis functions are step functions that are 1 within a bin and 0 elsewhere, so any sample is placed only in the bin that covers its position. In fact, a B-spline of order *k* places each sample into *k* bins, meaning that a B-spline estimator with *k* = 1 is equivalent to the Maximum Likelihood estimator. This is because there are *k* basis functions defined at each value of *z*, and each basis function corresponds to a single bin. Since the sum of the basis functions at any *z* is 1 we can use them to place a each point into *k* bins based on its *z*-value. This requires that we transform the expression values of a single gene to the range of the knot vector using

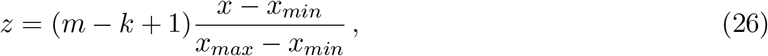

where *x* is the expression of a single gene and *x_max_* and *x_min_* are its maximum and minimum expression values.

Evaluating the B-spline for single gene gives a matrix *A*^x^ *∈ ℛ^N×m^* whose *ij*-th element is the weighting of sample *i* in bin *j*. We then compute the marginal probability of this gene using

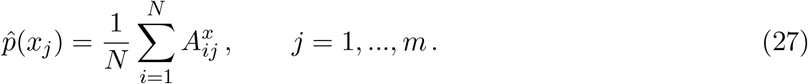

After obtaining the marginal distribution for another gene with expression profile *y* and obtaining *A^y^* we can compute the joint probability distribution using

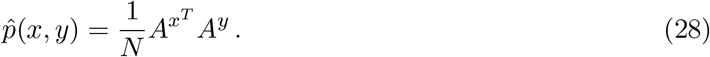

Then we compute the mutual information of *x* and *y* using (3)-(5).

#### S1.6 Kernel Density Estimator

Proposed by Moon et al. in [34], this estimator replaces rectangular histogram bins with kernels when estimating the probability distributions. The estimate of the probability density at expression *x* is given by

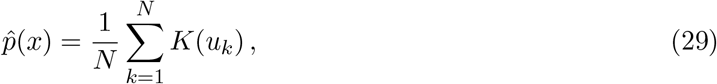

where *K*(*u_k_*) is the kernel function. This is a function of

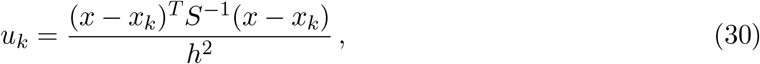

where *x_k_* is a measured expression value, *S* is the covariance matrix between the dimensions of *x_k_* and *h* is the kernel bandwidth. Note that for this estimator *x* is a vector of expression values at which we can sample the probability distribution and is not restricted to the measured expression values. In the original paper by Moon et al. the grid positions were chosen to be the locations of the data, but more recent implementations use a uniform grid over a range of [*x_min_ −* 1.5*h, x_max_* +1.5*h*], where *x_min_* and *x_max_* are the minimum and maximum measured expression values.

For marginal distributions *x* and *x_k_* are scalars and *S* is the variance of the expression profile whose density is being estimated. For joint distributions, *x* and *x_k_* are 2-D vectors and *S* is the 2x2 covariance matrix of the gene pair whose joint density is being estimated.

Using a *u_k_* of this form is known as the Fukunaga method [45], where the data has been linearly transformed to have a unit covariance matrix. This is equivalent to a whitening transformation of

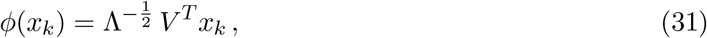

where Λ and *V* are the eigenvalue and eigenvector matrices of the sample covariance matrix of *x*.

To find the mutual information between two genes with expression profiles *x* and *y* the density estimates *p̂*(*x*), *p̂*(*y*) and *p̂*(*x, y*) sampled on the grid described above. The mutual information is then calculated using (2), where the sums are now over all grid positions.

This estimator has two parameters: the kernel function and the bandwidth. A common choice is the normalised Gaussian kernel:

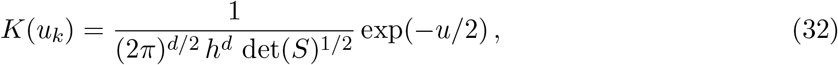

where *d* is the dimension of the density being estimated (1 for marginal distributions and 2 for joint distributions). The choice of *h* is more important than the choice of kernel when estimating probability densities. In the original paper, Moon et al. suggest using the optimal Gaussian bandwidth according to Silverman [46]:

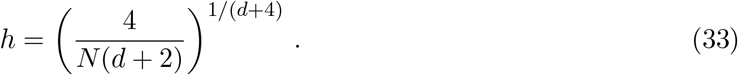

This choice minimises the mean squared error in *p̂*(*x*) if the true distribution is multivariate Gaussian and was found to give to comparable mutual information estimates to data-driven (and computationally expensive) bandwidth selection methods.

When performing density estimation for visualisation it is common to log-transform non-negative data to obtain a density estimate that is also non-negative. This should not affect the mutual information estimate due to the invariance of mutual information under homeomorphisms [35], but this study will investigate numerically the effect of the log-transformation on the mutual information estimate. It is also common to normalise the expression of each gene across all the samples to variance 1.

#### S1.7 *k*-Nearest-Neighbour Estimator

This approach utilises previous work by Kozachenko and Leonenko on estimating probability distributions and entropy from nearest neighbour distances [47], which was itself based on work by Vasicek [48]. Kraskov et al extended the analysis to produce a mutual information estimator [35].

For two expression profiles *x* and *y*, we make the pair *z* = (*x, y*). For this space we use the maximum norm

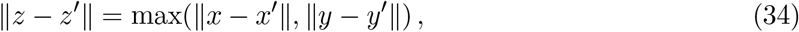

and compute *ε*(*i*)*/*2, the distance of *z_i_* = (*x_i_, y_i_*), *i* = 1, …, *N* to its *k*-th nearest neighbour. Using (34),

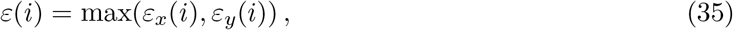

where *ε_x_*(*i*) and *ε_x_*(*i*) are the projections of *ε*(*i*) onto the *x* and *y* directions. We then count the number of samples with *x*∥*x_i_ − x_j_* ∥ < *ε_x_*(*i*)*/*2 for all other samples *j*, and label it *n_x_*(*i*). Similarly, we count the number of samples with ∥*y_i_ − y_j_* ∥ < *ε_y_* (*i*)*/*2 and label it *n_y_* (*i*). Finally, the estimate for the mutual information between *x* and *y* is

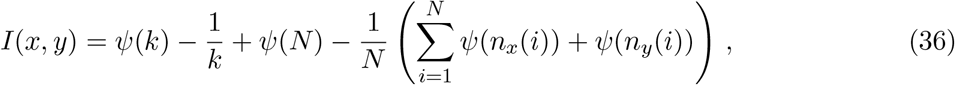

where *ψ*(*x*) is the digamma function satisfying

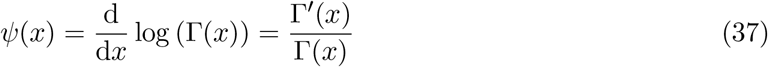

and Γ(*x*) is the gamma function.

This version of the *k*-NN estimator, also known as the KSG estimator after the authors of [35], has relatively few theoretical results but it has been shown that the KSG estimator struggles to detect stronger relationships [49]. This may make it inappropriate for gene network inference, where we are primarily concerned with detecting the strongest relationships between expression profiles.

Implementations of this estimator add a small random noise (*∼* 10^*−*9^) to break ties between nearest neighbour distances. The investigation presented in Section S5 found that the AUPRC only changes beyond its 5th decimal place for noise with maximum amplitudes of 10^*−*9^, 10^*−*10^ and 10^*−*11^.

#### S1.8 Parameters of the Histogram-based methods

Table 2 shows the parameters of each estimator. Further details on the parameters of the histogram-based estimators, the Maximum Likelihood, Miller-Madow, Chao-Shen and Shrinkage estimators is presented below.

##### S1.8.1 Binning method

For the Maximum Likelihood, Miller-Madow, Chao-Shen and Shrinkage estimators there are three binning methods available:

1. Equal width
2. Equal frequency
3. Bayesian Blocks

Histograms of the expression of the same gene are shown in Figure 9 using the three binning methods.

For equal width binning with *n* bins we divide the range of the data into bins with width (max *−* min)*/n*. Equal frequency binning uses *n* bins such that each has an equal frequency, which is equivalent to the area of the resulting histogram bar.

An alternative to these methods is the Bayesian Blocks algorithm, which was developed by Scargle et al. to model astronomical time series data using piecewise constant functions [16]. One application of the Bayesian Blocks algorithm is to place the times of discrete events into bins, where both the number and locations of the bins are chosen to best fit the data. This is equivalent to creating a histogram that is an optimal representation of data. This is achieved by optimising a fitness function, which can be any measure that quantifies how well a constant function fits the data in a bin. This work uses the default fitness function for this application,

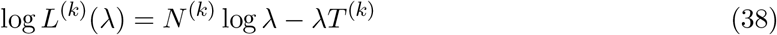

where *k* is the block index, *N* ^(*k*)^ is the number of counts in block *k*, *T* ^(*k*)^ is the length of the block and *λ* is the single parameter of the piecewise constant model.

The B-spline estimator can only be used with equal width bins.

### S2 Mutual Information Inference Algorithms

Three mutual information-based network inference algorithms will be used in this work. They are CLR, MRNET and ARACNE. As described in Section 2.1, each inference algorithm takes a symmetric matrix of mutual information value as an input. From this matrix the algorithm computes a list of scores, which are ranked in descending order. If a score between two genes has a high rank this reflects a confident prediction of an edge between them.

The implementations of the inference algorithms were taken from the R package *minet* [37].

The procedure by which the scores are computed is designed in order to remove indirect connections. Examples of network motifs that contain indirect connections are illustrated in Figure S1.

**Figure S1:**
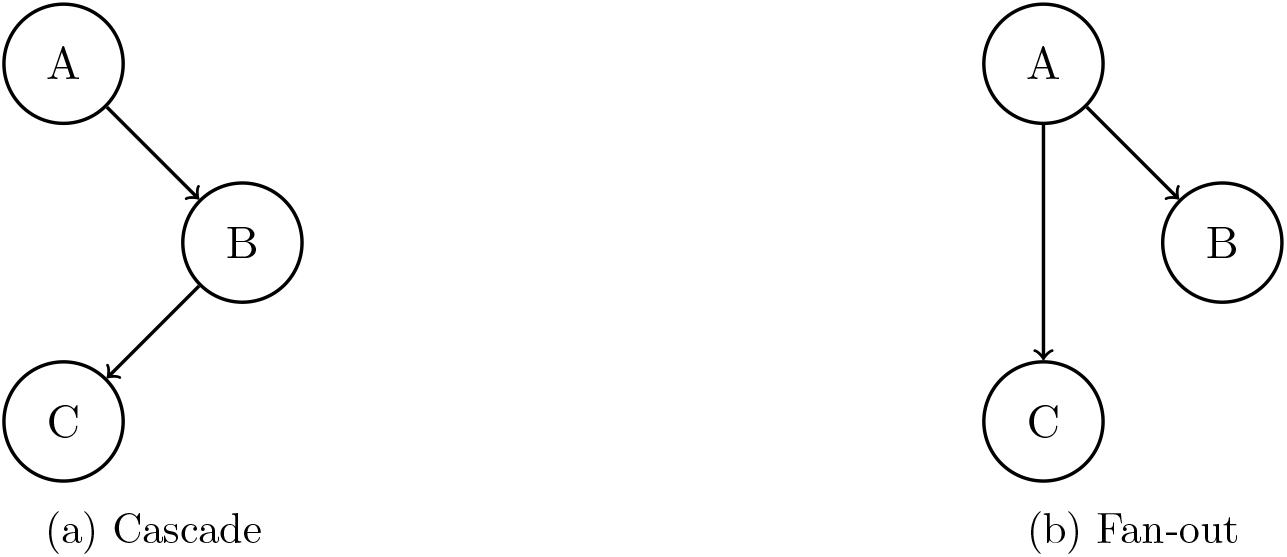
Two examples of network motifs that can cause systematic errors in network inference algorithms. This is due to dependence between the expression of two genes that are only indirectly connected in the true regulatory network. Network inference algorithms attempt to remove these indirect connections. In (a) there will be dependence between the expression of genes A and C, while in (b) there will be dependence between the expression of genes B and C.

#### S2.1 CLR

The Context Likelihood of Relatedness (CLR) algorithm is the most recent of the three inference algorithms and was develoed by Faith et al. in 2007 [9]. It considers the mutual information between the expression levels of a target gene and a second gene in the context of the distribution of the mutual information between the target gene and all other genes. The score of gene *i* with gene *j* is given by

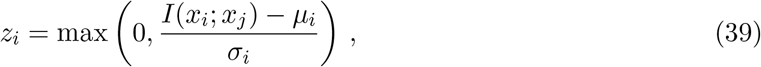

where *x_i_, x_j_* are the expressions of genes *i* and *j* and *μ_i_, σ_i_* are the mean and standard deviation of the distribution *I*(*x_i_, x_k_*), *k* = 1, …, *N, k ≠ i*. The final score between genes *i* and *j* is then given by 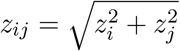.

#### S2.2 MRNET

MRNET is based on maximum relevance/minimum redundancy (MRMR), an information-theoretic feature selection technique that is common in machine learning [50]. “Maximum relevance” refers to choosing features that have a high mutual information with the target variable. “Minimum redundancy” means that these features are chosen such that the mutual information between them is as low as possible. Olsen et al. applied MRMR in a network inference context, in which the target is the expression of gene *i* and the features are the expression levels of all other genes. The steps of the algorithm are as follows:

1. Select the expression profile *x_i_* which has the largest mutual information with the target expression profile *y*.
2. Select the next expression profile *x_j_* as the one that maximizes

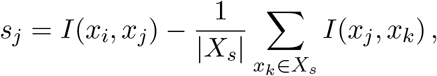

where *X_s_* is the set of previously selected expression profiles.
3. The score between genes *i* and *j* is given by *s_ij_* = max(*s_i_, s_j_*).

#### S2.3 ARACNE

The Algorithm for the Reconstruction of Accurate Cellular Networks was proposed by Margolin et al. in 2006 [6]. It starts by computing the mutual information between all the pairs of genes. These values are then filtered using a threshold that corresponds to a *p*-value in the null hypothesis of two independent genes. It then attempts to remove indirect connections using the Data Processing Inequality [51], which states that if an interaction between genes *i* and *j* depends on *k* then

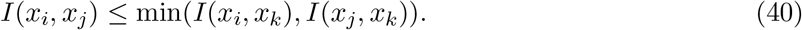

So for each gene triplet *i*, *j* and *k*, the lowest mutual information of *I*(*x_i_, x_j_*), *I*(*x_i_, x_k_*) and *I*(*x_j_, x_k_*) can only come from an indirect connection and so the corresponding edge is removed (its score is set to zero).

### S3 Data

This study will use the *in silico*, *e.coli* and *s.cerevisiae* datasets from the DREAM5 Network Inference Challenge [1]. The name *in silico* refers to the fact that this dataset has been simulated using GeneNetWeaver, a tool that uses ordinary differential equation models to simulate the expression of biologically inspired regulatory networks [18]. The *e.coli* and *s.cerevisiae* datasets use experimental data.

Each dataset consists of an expression matrix and a “gold standard” network, against which predictions are evaluated. The *in silico* expression matrix contains the raw values as simulated by GeneNetWeaver. The other two expression matrices have undergone microarray normalization followed by a log transform. Further details on the three datasets is displayed in Table 1.

The gold standard is only known for the *in silico* dataset. For the *e.coli* dataset was been constructed from RegulonDB, a database of experimentally verified transcriptional interactions [52]. The gold standard for the *s.cerevisiae* dataset was constructed following the reanalysis of ChIP-chip data [53]. The gold standards for the non-synthetic datasets are among the best in the field, but are almost certainly incomplete. Only interactions with strong experimental verification are included as edges in the network, so there may be many false negatives. Furthermore, only a subset of the genes are marked as potential regulators, and so the evaluation of an inferred network only occurs using edges between the regulators and other genes.

Detailed information on the source of all three datasets and the gold standard networks is available in the Supplementary Material of [1].

### S4 Evaluating network predictions-precision and recall

Network inference is a two-class classification task, where each pair of genes is classified as having an edge between them or not. Receiver operating characteristic (ROC) curves are a common method by which to evaluate such a classifier.

A predicted edge can be a true positive (TP), false positive (FP), true negative (TN) or false negative (FN). A ROC curve plots the true positive rate against the false positive rate

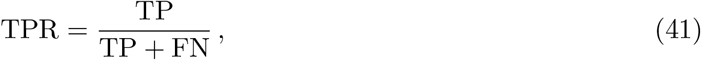

against the false positive rate

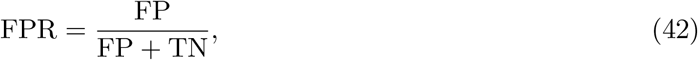

for various thresholds and evaluate the areas under the resulting curve (AUROC). A perfect classifier has area 1, while a random classifier has area 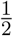 However, ROC curves are known to be potentially misleading in classification tasks with large class imbalances, which is the case in network inference as well as many other biological classification problems [19, 20]. Therefore we have a large number of true negatives that are of little interest when we evaluate our predicted networks.

As an alternative we use precision-recall curves, where

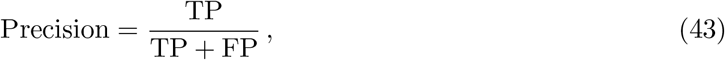

and Recall is equivalent to the true positive rate. Neither precision nor recall consider the number of true negatives. Similarly to a ROC curve, we plot the precision and recall for all thresholds. A more accurate classifier has a larger area under a precision-recall curve (AUPRC), which has a maximum value of 1 for a perfect classifier.

The F-or F1-score is twice the harmonic mean of the precision and recall and is given by

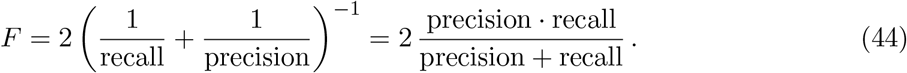

F-scores take values in [0, 1] and are close to 1 if both the precision and recall are close to 1, and close to 0 otherwise.

The area under precision recall curve was computed using the *PRROC* R package [54].

### S5 Finding the best parameters for each estimator

This section will identify the best parameters for the Maximum Likelihood, Miller-Madow, Chao-Shen, Shrinkage, B-spline and *k*-nearest-neighbours estimators. To recap, for the Maxiumum Like-lihood, Miller-Madow, Chao-Shen and Shrinkage estimators these are the number of bins and the binning method, unless the Bayesian Blocks binning method is used, in which case the number of bins are selected automatically. For the B-spline method we must choose the number of bins and the spline order. For the *k*-nearest-neighbour estimator we choose *k* and the amplitude of the noise added to each sample to break ties. The Kernel Density estimator has parameters (bandwidth and kernel function) but these were not investigated.

#### S5.1 *In silico* data

##### S5.1.1 Maximum Likelihood, Miller-Madow, Chao-Shen and Shrinkage estimators

First we compare the Maximum Likelihood, Miller-Madow, Chao-Shen and Shrinkage estimators. These estimators have been grouped together as they share the same parameters: binning method and number of bins. These results are shown in Figure S2. This work is not a comparison of inference algorithms, however it is worth noting that CLR (plots a, d, g and j in Figure S2) consistently has the largest AUPRC while ARACNE (plots c, f, i and l) has the smallest. This pattern is observed across all the mutual information estimators and datasets used in this study and will not be commented on again.

Using Bayesian Blocks increases the AUPRC for all 4 estimators when using CLR or ARACNE. For MRNET, Bayesian Blocks is the optimal parameter choice for the Miller-Madow and Shrinkage estimators.

For all of these estimators using *N* ^1*/*3^ bins is preferable to *N* ^1*/*2^ bins, but both of these choices give lower AUPRCs than Bayesian Blocks. Of these two choices, *N* ^1*/*3^ gives a larger AUPRC. This is probably due to *N* ^1*/*3^ being closer to the number of bins used by Bayesian Blocks than *N* ^1*/*2^x, as shown in Figure 5. The Freedman-Diaconis rule leads to a far lower AUPRC than the other choices of bin number for all the estimators and inference algorithms.

Since the Bayesian Blocks algorithm chooses the number of bins automatically, this improvement may be due to a more appropriate number of bins rather than the positioning of the bins. Plots g, h and i of Figure S2 show the AUPRC when using Equal Width and Equal Frequency bins with the number of bins chosen by Bayesian Blocks. These plots show that the locations of the bins also have a positive impact on the AUPRC, but that the majority of the increase in AUPRC when using Bayesian Blocks is from choosing an optimal number of bins.

From these results we can conclude that using Bayesian Blocks improves the performance of inference algorithms on this dataset. Furthermore, the number of bins is a more important parameter than the binning method for this dataset, in that if the number of bins is chosen “well,” then all the estimators are robust to the choice between equal frequency and equal width bins. Finally, using *N* ^1*/*2^ equal width bins with the shrinkage estimator results in a low AUPRC for all three inference algorithms.

##### S5.1.2 B-spline estimator

Figure S3 shows the AUPRC when using the B-spline estimator on the *in silico* dataset with various spline orders and bin numbers. For all three inference algorithms increasing the spline order generally increases the AUPRC, however for some choices of the number of bins this is not the case. For example, when using MRNET with 10 bins the AUPRC decreases as the spline order increases. For ARACNE the increase in AUPRC with spline order is smaller than for CLR or MRNET, and is barely noticeable.

Over the three inference algorithms and the various spline orders using *N* ^1*/*3^ gives either the largest or almost largest AUPRC. For this dataset the integer value of *N* ^1*/*3^ is 9, which is close to the recommended value of 10. Accordingly, the AUPRC between the two parameter choices are very similar. Using the same number of bins as Bayesian Blocks leads to a slightly lower AUPRC, and using *N* ^1*/*2^ bins is significantly worse. Once again, using the Freedman-Diaconis rule leads to a much lower AUPRC than any other parameter choice.

It is also interesting to note the increase in AUPRC for MRNET when using the B-spline estimator. When using the Maximum Likelihood, Miller-Madow, Chao-Shen and Shrinkage estimators the AUPRC of MRNET was significantly below CLR, however the difference is much smaller when using the B-spline estimator.

##### S5.1.3 *k*-Nearest-Neighbour estimator

Figure S4 shows the results for the *k*-NN estimator for a range of values of *k*. For both CLR and MRNET the AUPRC increases from *k* = 2 to *k* ≊ 10 and is approximately constant for larger values of *k*. For ARACNE this increase in AUPRC stops for *k ≥* 5. For CLR, MRNET and ARACNE the largest AUPRC is when *k* = 13, 14 and 14 respectively.

#### S5.2 *E. coli* data

##### S5.2.1 Maximum Likelihood, Miller-Madow, Chao-Shen and Shrinkage estimators

The AUPRC values for the *E. coli* dataset when using the Maximum Likelihood, Miller-Madow, Chao-Shen and Shrinkage estimators are shown in Figure S5. The AUPRC values are lower for this dataset than for the *in silico* dataset, which reflects the added difficulty of both inferring a network and obtaining an accurate gold standard for real biological systems. The variation in AUPRC between estimators, binning methods and bin numbers is also smaller for *E. coli* than for *in silico*.

Unlike for the *in silico* dataset, using Bayesian Blocks does not increase the AURPC for any of the mutual information estimator-inference algorithm combinations. However, the inference algorithms are more robust to the choice of bin number and binning method.

Equal width bins are preferable for all the estimators when using CLR and all but the Chao-Shen estimator for MRNET. For ARACNE, equal frequency bins give a larger AUPRC for the Maximum Likelihood, Miller-Madow and Shrinkage estimators.

For CLR, using *N* ^1*/*3^ bins (which are similar values, as shown by Figure 5) gives the highest AUPRC for all of these estimators. MRNET follows the same trend but the Chao-Shen estimator has the largest AUPRC when used with the same number of bins as Bayesian Blocks. For ARACNE, the Chao-Shen and Shrinkage estimators perform better when used with *N* ^1*/*2^ bins, while with the Maximum Likelihood and Chao-Shen estimators are best used with *N* ^1*/*3^ and the Bayesian Blocks number of bins respectively.

There are specific parameter choices that lead to low AUPRC values. The Freedman-Diaconis rule leads to the lowest AUPRC. However, this difference is negligible when using CLR or ARACNE with the Chao-Shen and Shrinkage estimators (plots j and l in Figure S5).

##### S5.2.2 B-spline estimator

Figure S6 shows the AUPRC when using the B-spline estimator on the *E. coli* dataset with various spline orders and bin numbers. Unlike for the *in silico* dataset, the AUPRC does not increase with spline order. When varying the number of bins the change in AUPRC is *∼* 0.001, except when using MRNET with the Freedman-Diaconis rule. In this case the AUPRC is significantly lower than all choices of bin number.

Yet again, CLR performs best with ARACNE performing worst, however, now the Freedman-Diaconis rule only leads to low AUPRC values when using MRNET. For both CLR and ARACNE the number of bins does not strongly affect the AUPRC.

##### S5.2.3 *k*-Nearest-Neighbour Estimator

The same trends are apparent for the *E. coli* data as for the *in silico* dataset, however the difference between low and high values of *k* for CLR and MRNET are now smaller than for the *in silico* data.

#### S5.3 S. cerevisiae data

##### S5.3.1 All estimators

The AUPRC values for the various estimators are shown in Figures S8, S9 and S10. This dataset has the lowest AUPRC values andn there is very little variation in AUPRC between different parameters or estimators. The variation between inference algorithms is also lower than for the *in silico* and *E. coli* datasets.

### S6 Best estimator parameters by inference algorithm-MRNET and ARACNE

Tables S1 and S2 show the estimator parameters that maximised the AUPRC when using MRNET and ARACNE respectively.

**Table S1:**
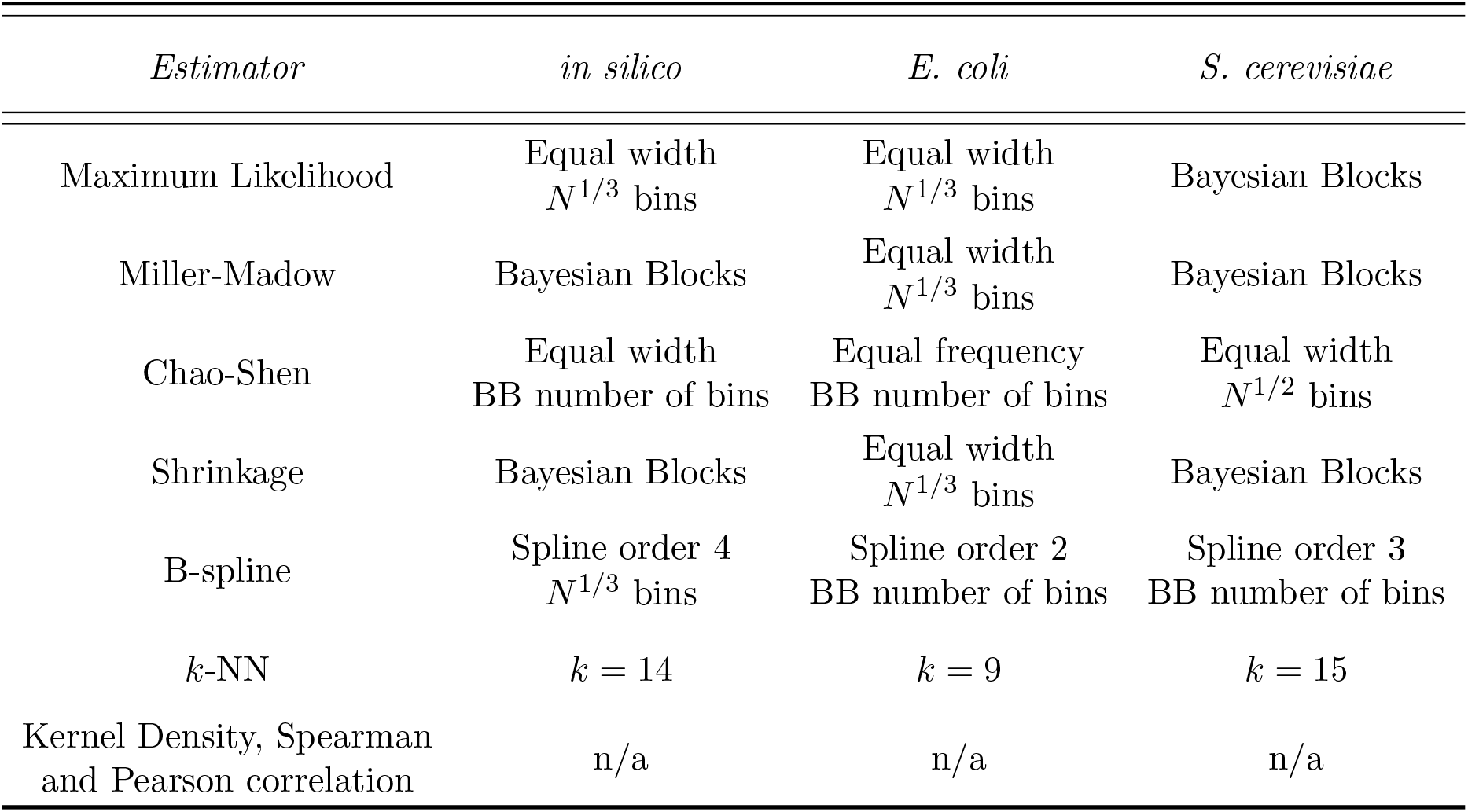
The mutual information estimator parameters that maximise the AUPRC when using the MRNET network inference algorithm. The parameters of each estimator are shown in Table 2 and described in detail in Section S1. The parameters of the Kernel Density estimator were not investigated and the only the values suggested by the authors of [34] were used. The Spearman and Pearson correlation estimators do not have parameters.

| *Estimator* | *in silico* | *E. coli* | *S. cerevisiae* |
| --- | --- | --- | --- |
| Maximum Likelihood | Equal width<br>$N^{1/3}$ bins | Equal width<br>$N^{1/3}$ bins | Bayesian Blocks |
| Miller-Madow | Bayesian Blocks | Equal width<br>$N^{1/3}$ bins | Bayesian Blocks |
| Chao-Shen | Equal width<br>BB number of bins | Equal frequency<br>BB number of bins | Equal width<br>$N^{1/2}$ bins |
| Shrinkage | Bayesian Blocks | Equal width<br>$N^{1/3}$ bins | Bayesian Blocks |
| B-spline | Spline order 4<br>$N^{1/3}$ bins | Spline order 2<br>BB number of bins | Spline order 3<br>BB number of bins |
| $k$-NN | $k = 14$ | $k = 9$ | $k = 15$ |
| Kernel Density, Spearman<br>and Pearson correlation | n/a | n/a | n/a |

**Table S2:** The mutual information estimator parameters that maximise the AUPRC when using the ARACNE network inference algorithm. The parameters of each estimator are shown in Table 2 and described in detail in Section S1. The parameters of the Kernel Density estimator were not investigated and the only the values suggested by the authors of [34] were used. The Spearman and Pearson correlation estimators do not have parameters.

| *Estimator* | *in silico* | *E. coli* | *S. cerevisiae* |
| --- | --- | --- | --- |
| Maximum Likelihood | Bayesian Blocks | Equal frequency<br>$N^{1/3}$ bins | Bayesian Blocks |
| Miller-Madow | Bayesian Blocks | Equal frequency<br>BB number of bins | Bayesian Blocks |
| Chao-Shen | Bayesian Blocks | Equal width<br>$N^{1/2}$ bins | Bayesian Blocks |
| Shrinkage | Bayesian Blocks | Equal frequency<br>$N^{1/2}$ bins | Equal width<br>BB number of bins |
| B-spline | Spline order 5<br>10 bins | Spline order 4<br>$N^{1/2}$ bins | Spline order 3<br>FD number of bins |
| $k$-NN | $k = 14$ | $k = 3$ | $k = 15$ |
| Kernel Density, Spearman<br>and Pearson correlation | n/a | n/a | n/a |

**Figure S2:**
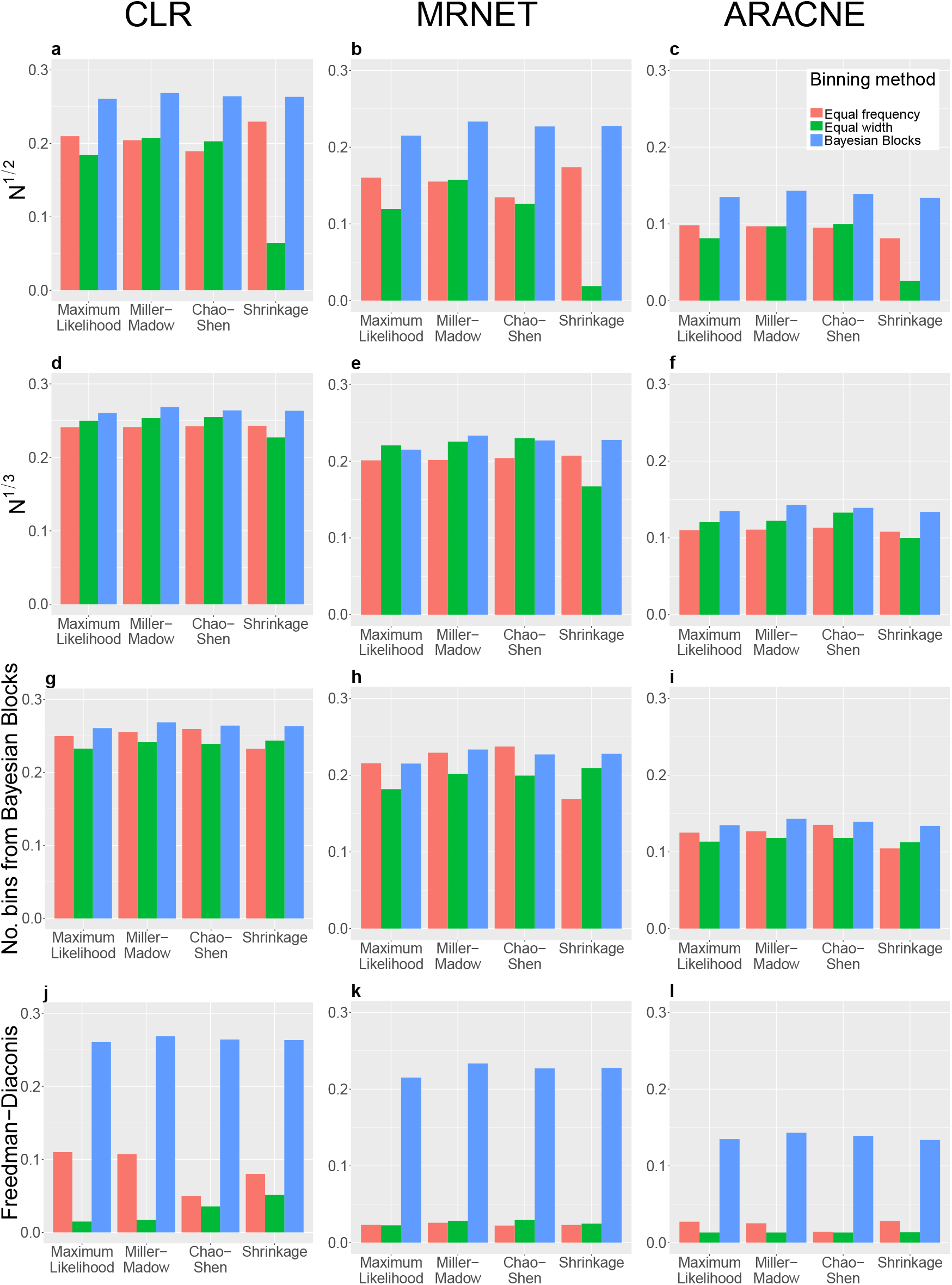
The AUPRC for the *in silico* dataset with Maximum Likelihood, Miller-Madow, Chao-Shen and Shrinkage estimators. An individual plot shows the AUPRC when using a single inference algorithm and a specific number of bins with the four estimators. Each grouping of bars represents a single estimator for the three binning methods. Each column shows results for a single inference algorithm and each row shows results for a single number of bins. Note that when using the Bayesian Blocks binning method the number of bins is chosen automatically, hence the AUPRC values within columns are the same for the same MI estimator, but are included in all plots for ease of comparison.

**Figure S3:**
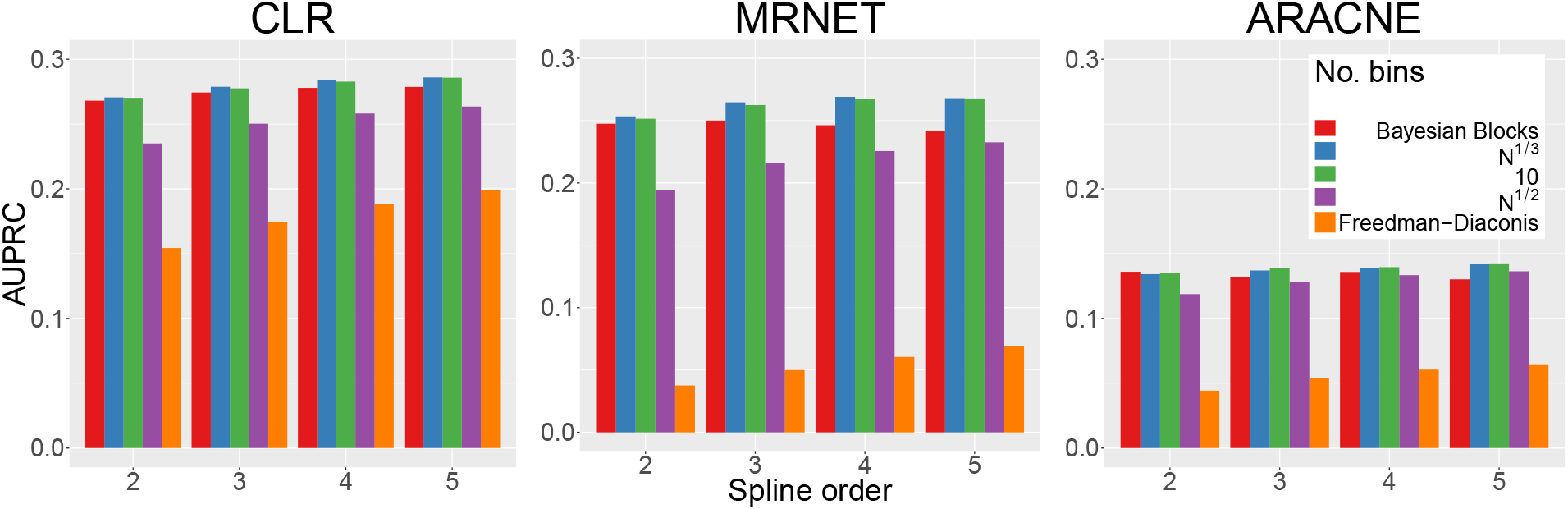
The AUPRC results for the *in silico* dataset with the B-spline estimator. Each plot shows the AUPRC results for a single inference algorithm, with the results being grouped by spline order.

**Figure S4:**
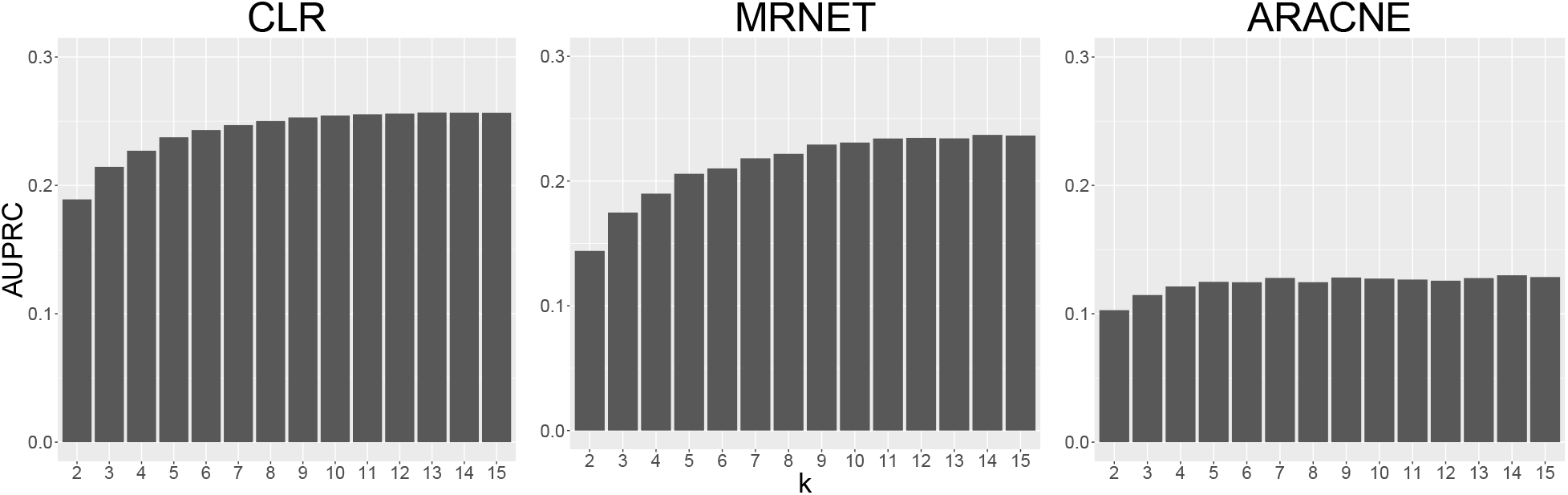
The AUPRC results when using the *k*-Nearest-Neighbour estimator on the *in silico* dataset. Each plot contains the AUPRC results for a single inference algorithm and each bar represents the AUPRC when using *k* nearest neighbours to estimate the mutual information.

**Figure S5:**
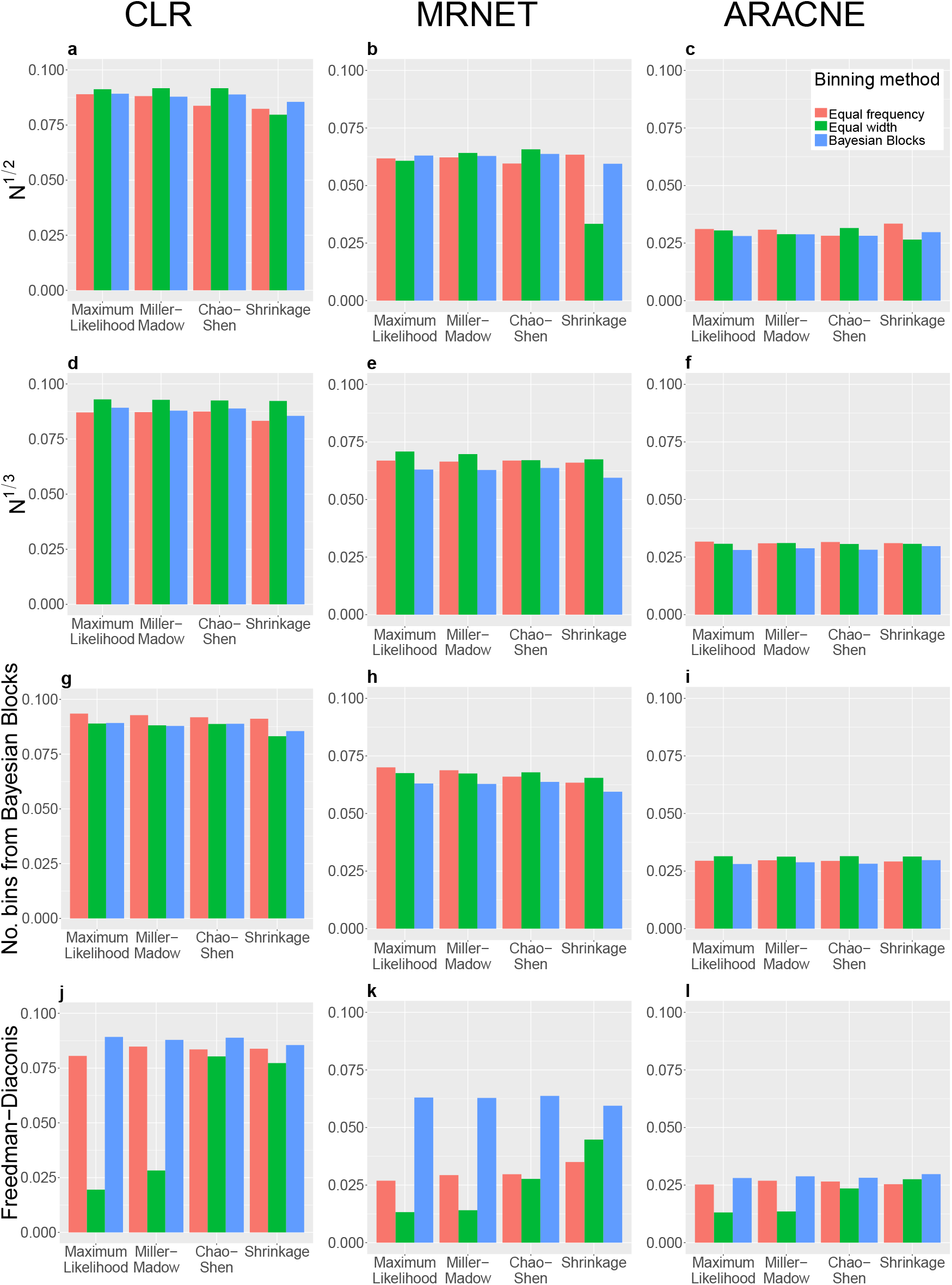
The AUPRC for the *E. coli* dataset with Maximum Likelihood, Miller-Madow, Chao-Shen and Shrinkage estimators. An individual plot shows the AUPRC when using a single inference algorithm and a specific number of bins with the four estimators. Each grouping of bars represents a single estimator for the three binning methods. Each column shows results for a single inference algorithm and each row shows results for a single number of bins. Note that when using the Bayesian Blocks binning method the number of bins is chosen automatically, hence the AUPRC values within columns are the same for the same MI estimator, but are included in all plots for ease of comparison.

**Figure S6:**
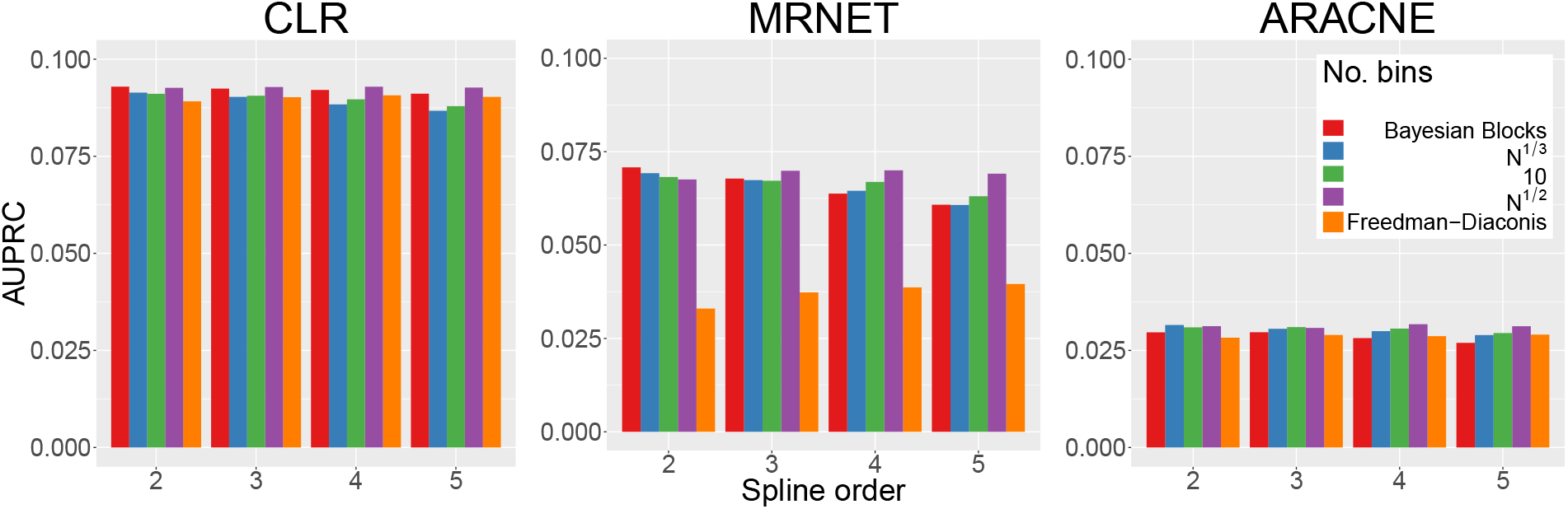
The AUPRC results for the *E. coli* dataset with the B-spline estimator. Each plot shows the AUPRC results for a single inference algorithm, with the results being grouped by spline order.

**Figure S7:**
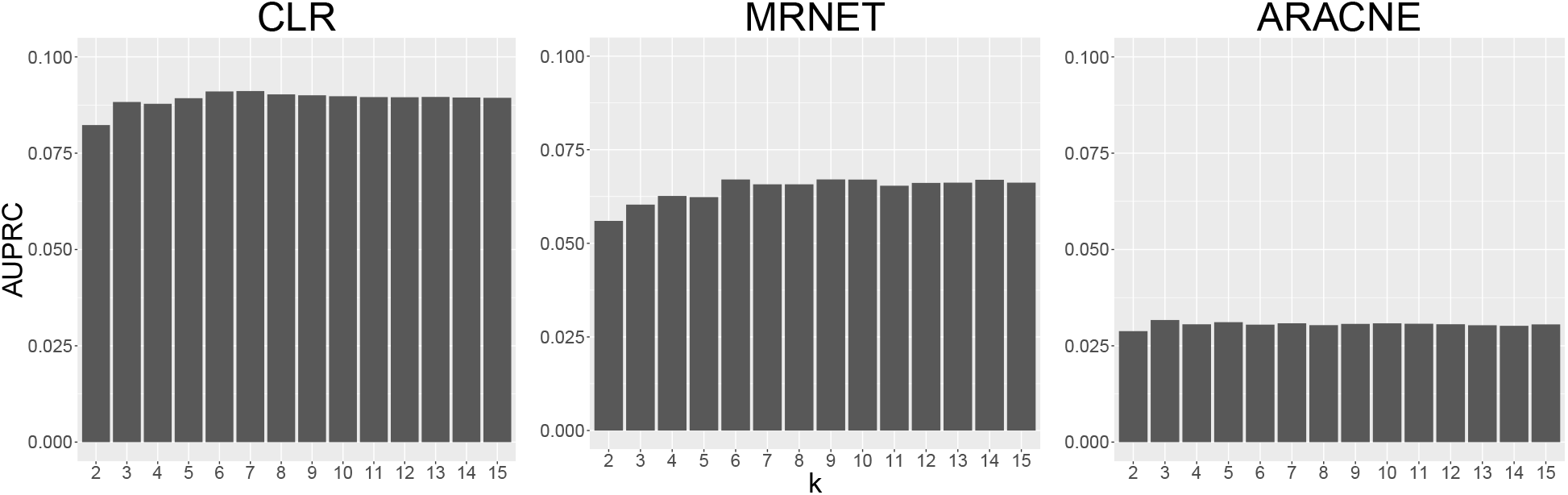
The AUPRC results when using the *k*-Nearest-Neighbour estimator on the *E. coli* dataset. Each plot contains the AUPRC results for a single inference algorithm and each bar represents the AUPRC when using *k* nearest neighbours to estimate the mutual information.

**Figure S8:**
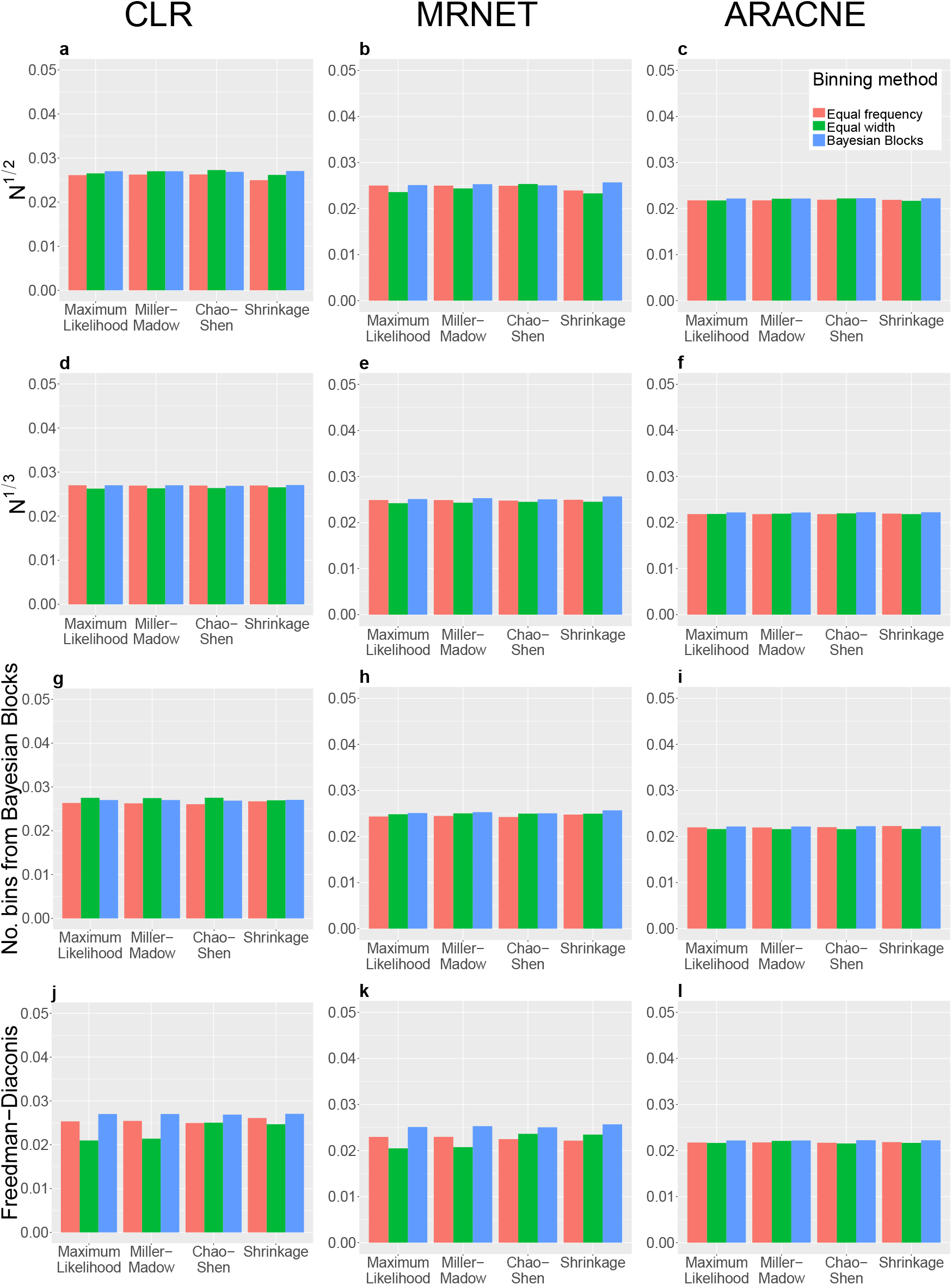
The AUPRC for the *S. cerevisiae* dataset with Maximum Likelihood, Miller-Madow, Chao-Shen and Shrinkage estimators. An individual plot shows the AUPRC when using a single inference algorithm and a specific number of bins with the four estimators. Each grouping of bars represents a single estimator for the three binning methods. Each column shows results for a single inference algorithm and each row shows results for a single number of bins. Note that when using the Bayesian Blocks binning method the number of bins is chosen automatically, hence the AUPRC values within columns are the same for the same MI estimator, but are included in all plots for ease of comparison.

**Figure S9:**
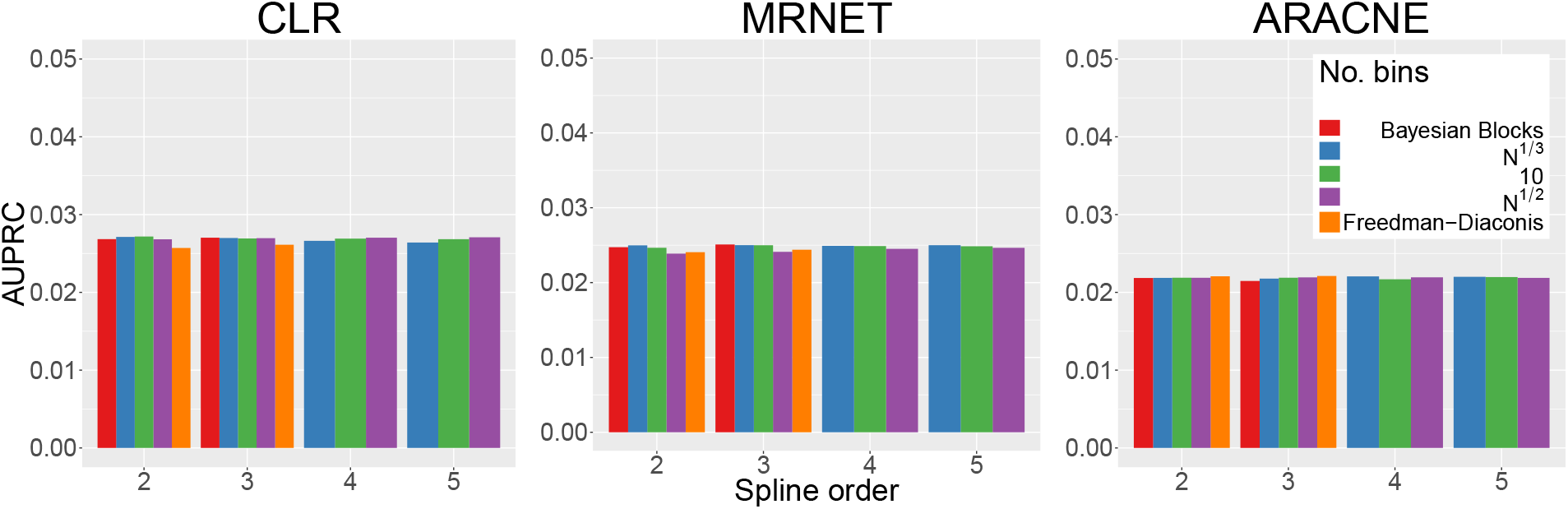
The AUPRC results for the *S. cerevisiae* dataset with the B-spline estimator. Each plot shows the AUPRC results for a single inference algorithm, with the results being grouped by spline order.

**Figure S10:**
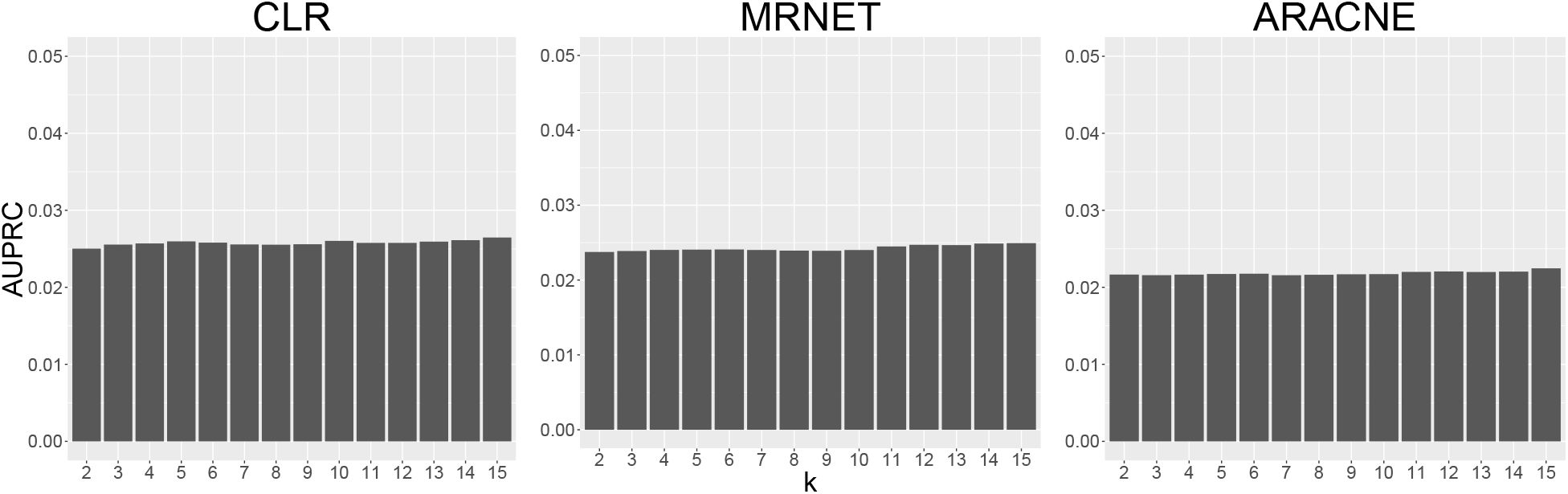
The AUPRC results when using the *k*-Nearest-Neighbour estimator on the *S. cerevisiae* dataset. Each plot contains the AUPRC results for a single inference algorithm and each bar represents the AUPRC when using *k* nearest neighbours to estimate the mutual information.

